# Genomic Stability and Genetic Defense Systems in *Dolosigranulum pigrum* a Candidate Beneficial Bacterium from the Human Microbiome

**DOI:** 10.1101/2021.04.16.440249

**Authors:** Stephany Flores Ramos, Silvio D. Brugger, Isabel Fernandez Escapa, Chelsey A. Skeete, Sean L. Cotton, Sara M. Eslami, Wei Gao, Lindsey Bomar, Tommy H. Tran, Dakota S. Jones, Samuel Minot, Richard J. Roberts, Christopher D. Johnston, Katherine P. Lemon

**Affiliations:** The Forsyth Institute (Microbiology), Cambridge, MA, USA; Department of Infectious Diseases and Hospital Epidemiology, University Hospital Zurich, University of Zurich, Zurich, Switzerland; Department of Oral Medicine, Infection and Immunity, Harvard School of Dental Medicine, Boston, MA, USA; Alkek Center for Metagenomics & Microbiome Research, Department of Molecular Virology & Microbiology, Baylor College of Medicine, Houston, Texas, USA; Vaccine and Infectious Diseases Division, Fred Hutchinson Cancer Research Center, Seattle, WA, USA; New England Biolabs, Ipswich, MA, USA; Division of Infectious Diseases, Boston Children’s Hospital, Harvard Medical School, Boston, MA, USA; Section of Infectious Diseases, Texas Children’s Hospital, Department of Pediatrics, Baylor College of Medicine, Houston, Texas, USA

**Keywords:** *Dolosigranulum pigrum*, nasal microbiota, pangenome, methylome, restriction modification, CRISPR, mobile genetic elements

## Abstract

*Dolosigranulum pigrum* is positively associated with indicators of health in multiple epidemiological studies of human nasal microbiota. Knowledge of the basic biology of *D. pigrum* is a prerequisite for evaluating its potential for future therapeutic use; however, such data are very limited. To gain insight into *D. pigrum*’s chromosomal structure, pangenome and genomic stability, we compared the genomes of 28 *D. pigrum* strains that were collected across 20 years. Phylogenomic analysis showed closely related strains circulating over this period and closure of 19 genomes revealed highly conserved chromosomal synteny. Gene clusters involved in the mobilome and in defense against mobile genetic elements (MGEs) were enriched in the accessory genome versus the core genome. A systematic analysis for MGEs identified the first candidate *D. pigrum* prophage and insertion sequence. A systematic analysis for genetic elements that limit the spread of MGEs, including restriction modification (RM), CRISPR-Cas, and deity-named defense systems, revealed strain-level diversity in host defense systems that localized to specific genomic sites including one RM system hotspot. Analysis of CRISPR spacers pointed to a wealth of MGEs against which *D. pigrum* defends itself. These results reveal a role for horizontal gene transfer and mobile genetic elements in strain diversification while highlighting that in *D. pigrum* this occurs within the context of a highly stable chromosomal organization protected by a variety of defense mechanisms.

**IMPORTANCE:** *Dolosigranulum pigrum* is a candidate beneficial bacterium with potential for future therapeutic use. This is based on its positive associations with characteristics of health in multiple studies of human nasal microbiota across the span of human life. For example, high levels of *D. pigrum* nasal colonization in adults predicts the absence of *Staphylococcus aureus* nasal colonization. Also, *D. pigrum* nasal colonization in young children is associated with healthy control groups in studies of middle ear infections. Our analysis of 28 genomes revealed a remarkable stability of *D. pigrum* strains colonizing people in the U.S. across a 20-year span. We subsequently identified factors that can influence this stability, including genomic stability, phage predators, the role of MGEs in strain-level variation and defenses against MGEs. Finally, these *D. pigrum* strains also lacked predicted virulence factors. Overall, these findings add additional support to the potential for *D. pigrum* as a therapeutic bacterium.

## INTRODUCTION

Evidence points to a prominent role for the benign nasal bacterium *Dolosigranulum pigrum* in structuring nasal microbiota beneficial to human health (1–30) (reviewed in (31–36)). Individuals whose nasal microbiota is dominated by *D. pigrum* are less likely to be colonized by nasal pathobionts and, therefore, are at lower risk of invasive infections due to these microbes. For example, *D. pigrum* is inversely associated with *Staphylococcus aureus* in adult nostrils (5, 16, 28, 37). Also, the level of maternal *D. pigrum* is inversely associated with infant acquisition of *S. aureus* (38), and, in a small study, neonates who do not acquire *S. aureus* have a higher relative abundance of *D. pigrum* (39). During *in vitro* growth, *D. pigrum* inhibits *S. aureus* on agar medium, but not the reverse (28), suggesting *D. pigrum* might directly antagonize *S. aureus in vivo*. Additionally, *D. pigrum* and nasal *Corynebacterium* species are frequently present in pediatric nasal microbiota when *Streptococcus pneumoniae* is absent (1, 8). Together *D. pigrum* and *Corynebacterium pseudodiphtheriticum* robustly inhibit *S. pneumoniae in vitro*, as compared to either alone (28). As illustrated by these examples, nasal microbiota with higher levels of *D. pigrum*—usually alongside *Corynebacterium*—are often associated with health. Young infants with prolonged high levels of *D. pigrum* and *Corynebacterium* exhibit greater stability of their nasal microbiota and fewer respiratory tract infections (3, 4, 6, 11, 21). Also, higher levels of nasal *D. pigrum* and *Corynebacterium* are more common in healthy children than in those with pneumonia (12) or those with otitis media (1, 2, 15, 30).

In stark contrast to the steadily increasing data in support of *D. pigrum* as a candidate beneficial bacterium (40), there is a dearth of information about the basic biology of this Gram-positive organism, including the organization and stability of its genome. Ideally, bacterial strains with therapeutic potential display a reliably stable genome structure and have the capacity to resist horizontal gene transfer (HGT), since the latter might lead to unanticipated effects. The stability of bacterial genomes reflects a balance between competing factors including invasion by mobile genetic elements (MGEs) and systems that defend against these. MGEs play a key role in strain variation through acquisition and distribution of genes in the accessory genome. Analysis of the pangenome of multiple strains identifies core and soft-core gene clusters (GCs) common to all, or almost all, of the strains, respectively, and GCs present in smaller subsets of strains, which constitute the accessory genome (41, 42). Although accessory genes may result from gene loss, many are thought to be acquired via HGT. Counterbalancing this are key systems for defense against MGEs. These include well-described restriction modification systems, CRISPR-Cas systems, and the more recently identified, deity-named defense systems (43). RM systems distinguish intracellular DNA as self or nonself by virtue of specific methyl-modifications within short linear sequences that allow for destruction of inappropriately methylated nonself DNA by endonuclease activity; the various RM systems are classified into Type I, II, III and IV. There are also other variations of DNA modification-based defense (44, 45). CRISPR-Cas systems mediate defense using a multistep process. Small fragments of foreign nucleic acids are first recognized as non-self and incorporated into the host genome between short DNA repeats, known as a CRISPR array. Subsequently, these fragments, now spacers within the array, are used as RNA guiding molecules for an endonuclease complex that recognizes and destroys DNA containing these sequences (46). The more recently identified deity-named defense systems consist of a set of ten disparate antiphage/plasmid mechanisms that are often found clustered next to known defense genes (RM and CRISPR-Cas) (43) within defense islands (47) of bacterial genomes. Although deity-named defense systems have been shown to be active and limit phage/plasmid spread, their exact underlying modes of action remain to be deciphered. Collectively, these systems can protect bacteria from infection by phages and invasion by other MGEs, including plasmids and transposable elements, thus limiting introduction of new genes and maintaining genomic stability.

Comparing genomic content and chromosomal organization of *D. pigrum* strains collected 20 years apart, and mostly in the U.S., we identified the following characteristics: 1) highly similar strains circulating across 20 years; 2) stable chromosomal synteny across the phylogeny; 3) the first predicted *D. pigrum* prophage and insertion sequence; and 4) a diverse collection of RM, deity-named defense and CRISPR-Cas systems incorporated at conserved chromosomal insertion sites across strains. Together, these reveal a stable synteny and a high-level of sequence conservation within the *D. pigrum* core genome, along with an open pangenome and active defense against HGT.

## RESULTS

### Detection of highly similar *Dolosigranulum pigrum* strains over a 20-year span

To identify genomic shifts in *D. pigrum* strains currently circulating in human nasal microbiota compared to strains from approximately 20 years ago, we collected 17 new nostril isolates of *D. pigrum* from volunteers in 2017-2018 and sequenced the genomes of these isolates using SMRTSeq (PacBio), fully circularizing 15 (**Table 1**). We compared these 17 new genomes to 11 described genomes (28), 9 of which are from strains collected in the late 1990s (48). This refined existing and uncovered new information about the basic genomic characteristics of *D. pigrum* (**File S1, Table A**).

**Table 1.** Source information for the 28 *D. pigrum* strains and Quality description for the 17 newly SMART-sequenced closed genomes

| Original strain name | Internal reference | Year Isolated | Human body site | Geo-graphy | Age (years) | NCBI assembly ID | Citation | Realigned Bases %<sup>#</sup> | Coverage (mean) |
| --- | --- | --- | --- | --- | --- | --- | --- | --- | --- |
| ATCC 51524 | n.a. | 1988 | spinal cord | UK | ? | GCF_000245815.1 | Aguirre 1993 |  |  |
| KPL1914 | KPL1914 | 2010 | nostril | MA | Adult | GCA_003263915.2 | Brugger 2020 |  |  |
| CDC 39-95 | KPL1922 | 1995 | np | CN | 3 | GCF_003264145.1 | LaClaire 2000 |  |  |
| CDC 2949-98 | KPL1930 | 1998 | np | AZ | ? | GCF_003264135.1 | LaClaire 2000 |  |  |
| CDC 4294-98 | KPL1931 | 1998 | blood | SC | < 1 | GCF_003264085.1 | LaClaire 2000 |  |  |
| CDC 4420-98 | KPL1932 | 1998 | blood | TN | 11 | GCF_003264065.1 | LaClaire 2000 |  |  |
| CDC 4545-98 | KPL1933 | 1998 | np | AZ | ? | GCF_003264045.1 | LaClaire 2000 |  |  |
| CDC 4709-98 | KPL1934 | 1998 | eye | GA | < 1 | GCA_003264015.2 | LaClaire 2000 |  |  |
| CDC 4199-99 | KPL1937 | 1999 | blood | GA | ~ 2 | GCF_003264005.1 | LaClaire 2000 |  |  |
| CDC 4791-99 | KPL1938 | 1999 | np | AZ | ? | GCF_003263975.1 | LaClaire 2000 |  |  |
| CDC 4792-99 | KPL1939 | 1999 | np | AZ | ? | GCF_003263965.1 | LaClaire 2000 |  |  |
| KPL3033 | KPL3033 | 2018 | nostril | MA | 18-30 | GCA_017655925.1 | this study | 92.61% * | 498 |
| KPL3043 | KPL3043 | 2018 | nostril | MA | 7-12 | GCA_017655905.1 | this study | 92.40% * | 582 |
| KPL3050 | KPL3050 | 2018 | nostril | MA | 31-60 | GCA_017655885.1 | this study | 92.11% * | 475 |
| KPL3052 | KPL3052 | 2018 | nostril | MA | 3-6 | GCA_017655865.1 | this study | 92.15% * | 382 |
| KPL3065 | KPL3065 | 2018 | nostril | MA | 7-12 | GCA_017655845.1 | this study | 91.73% * | 460 |
| KPL3069 | KPL3069 | 2018 | nostril | MA | 7-12 | GCA_017655825.1 | this study | 88.13% * | 372 |
| KPL3070 | KPL3070 | 2018 | nostril | MA | 31-60 | GCA_017655785.1 | this study | 91.85% * | 271 |
| KPL3077 | KPL3077 | 2018 | nostril | MA | 7-12 | GCA_017655765.1 | this study | 91.60% | 351 |
| KPL3084 | KPL3084 | 2018 | nostril | MA | 31-60 | GCA_017655745.1 | this study | 90.24% * | 433 |
| KPL3086 | KPL3086 | 2018 | nostril | MA | < 3 | GCA_017655725.1 | this study | 91.30% * | 342 |
| KPL3090 | KPL3090 | 2018 | nostril | MA | 7-12 | GCA_017655685.1 | this study | 90.72% * | 423 |
| KPL3246 | KPL3246 | 2018 | nostril | MA | 7-12 | GCA_017655805.1 | this study | 92.47% * | 578 |
| KPL3250 | KPL3250 | 2018 | nostril | MA | 7-12 | GCA_017655665.1 | this study | 92.63% * | 501 |
| KPL3256 | KPL3256 | 2018 | nostril | MA | 7-12 | GCA_017655645.1 | this study | 92.84% | 530 |
| KPL3264 | KPL3264 | 2018 | nostril | MA | 7-12 | GCA_017655705.1 | this study | 87.61% | 342 |
| KPL3274 | KPL3274 | 2018 | nostril | MA | 7-12 | GCA_017655945.1 | this study | 87.41% * | 574 |
| KPL3911 | KPL3911 | 2017 | nostril | MA | < 3 | GCA_017655965.1 | this study | 87.13% * | 595 |

<sup>#</sup> Percent Realigned Bases (from Realignment to Draft Assembly). \* Circularized genome

To assess the similarity of these 28 *D. pigrum* strains, we generated a phylogenomic tree based on single-copy core GCs (**Fig. 1**). This phylogeny revealed a relatively small evolutionary distance among *D. pigrum* strains. The average number of pairwise single nucleotide polymorphisms (SNPs) among isolates collected approximately 20 years apart was similar to that among isolates collected recently (21,754 vs. 20,834) (**Table S1**). Thus, closely related strains of *D. pigrum* have circulated among people in the U.S. over a span of time that has an upper bound of 20 years and a lower bound of 8-13 years. (This lower bound allows for the possibility that the recent isolates were stably acquired in infancy since most of the 2018 strains were from children in the 7-12-year age range.) In contrast to the *D. pigrum*-only phylogeny (**Fig. 1**), a phylogeny using *Alloiococcus otitis* (49) (**File S1, Fig. A**), the closest genome-sequenced bacterium to *D. pigrum* in 16S rRNA gene phylogenies, as an outgroup displayed poor node support, likely due to poor SNP resolution (**Table S1**). Therefore, we based subsequent inferences on the *D. pigrum*- only phylogeny.

**Figure 1.**
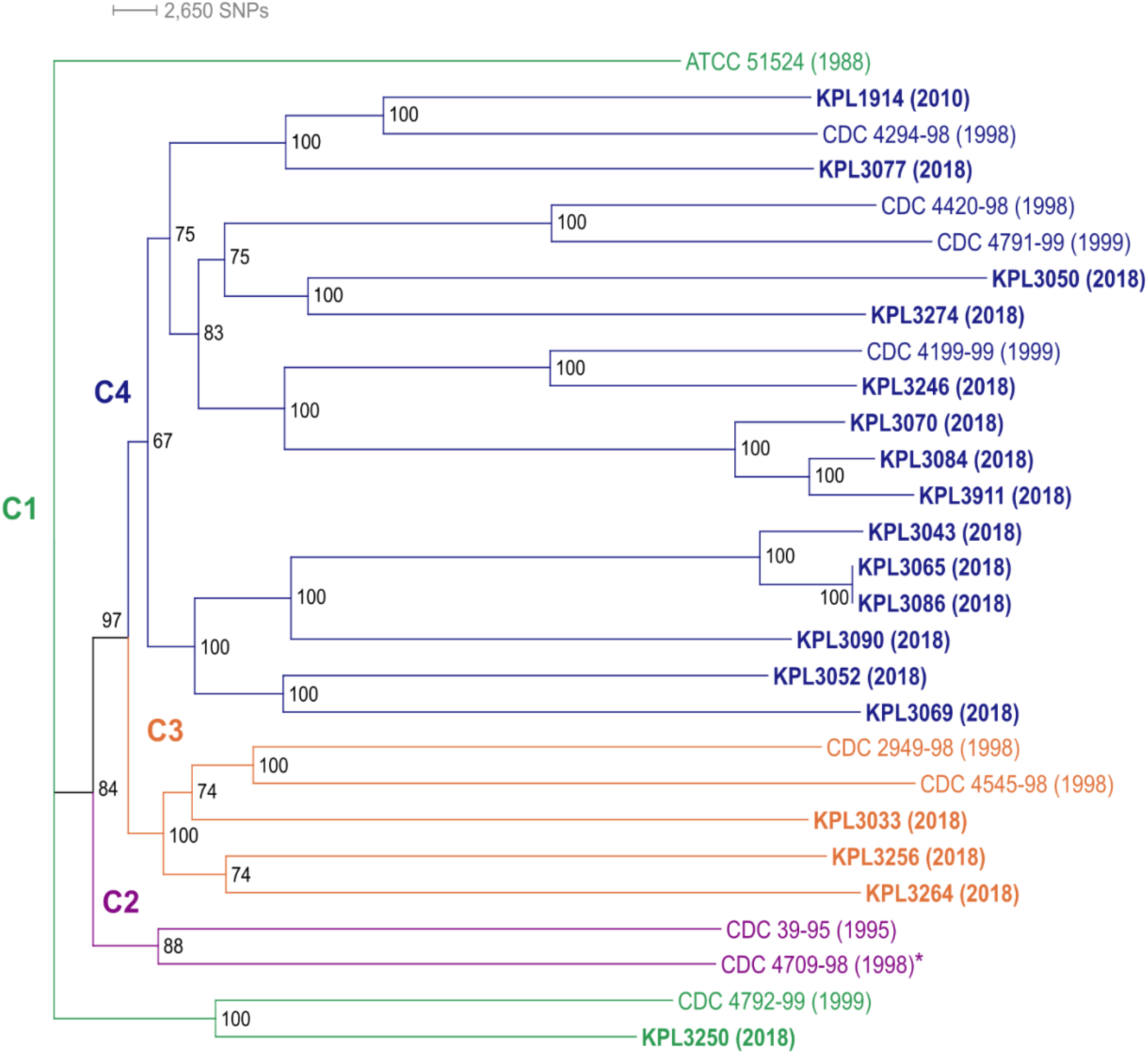
*Dolosigranulum pigrum* strains collected 20 years apart are phylogenetically similar. This maximum likelihood core-gene-based phylogeny shows recently collected strains (bold), mostly from 2018, and strains collected before 2000 intermingled in three of the four distinct clades (clades C1-C4 are color coded, year of collection is in parentheses). Strains separated by 12 to 20 years grouped together in terminal clades: KPL1914 and CDC 4294-98; KPL3246 and CDC 4199-99; and KPL3250 with CDC 4792-99. Genomes of strains in bold plus strain CDC 4709-98 (asterisk) are closed. Strains KPL3065 and KPL3086 were collected from two different individuals and have almost identical genomes, differing by just 4 core SNPs and 6 gene clusters (4 and 2 in KPL3086 and KPL3065, respectively). We created this unrooted phylogeny using the concatenated alignment of 1102 conservative single-copy core GCs (**File S1, Figure Bi**), a GTR+F+R3 substitution model of evolution, 553 maximum likelihood searches, and 1000 ultrafast Bootstraps with IQ-Tree v. 1.

### The chromosome of *D. pigrum* exhibits conserved synteny across a phylogeny spanning 20 years

Based on the observed similarity of circulating strains over time, we hypothesized there would be a high-level of genomic stability across the *D. pigrum* phylogeny. To test this, we compared chromosomal synteny across the four major clades in the *D. pigrum* phylogeny using 19 strains with closed genome sequences (highlighted in bold or with * in **Fig. 1**), including representative strains collected in 1998, 2010, 2017, and 2018. A MAUVE alignment (50, 51) of these 19 genomes starting at the *dnaA* gene revealed a remarkable conservation of the overall chromosomal structure with no visible shifts in the position of large blocks of sequence (**Fig. 2A**). Dispersed among these blocks are regions with higher numbers of insertions and deletions (indels) (**Fig. 2A; File S2**).

**Figure 2.**
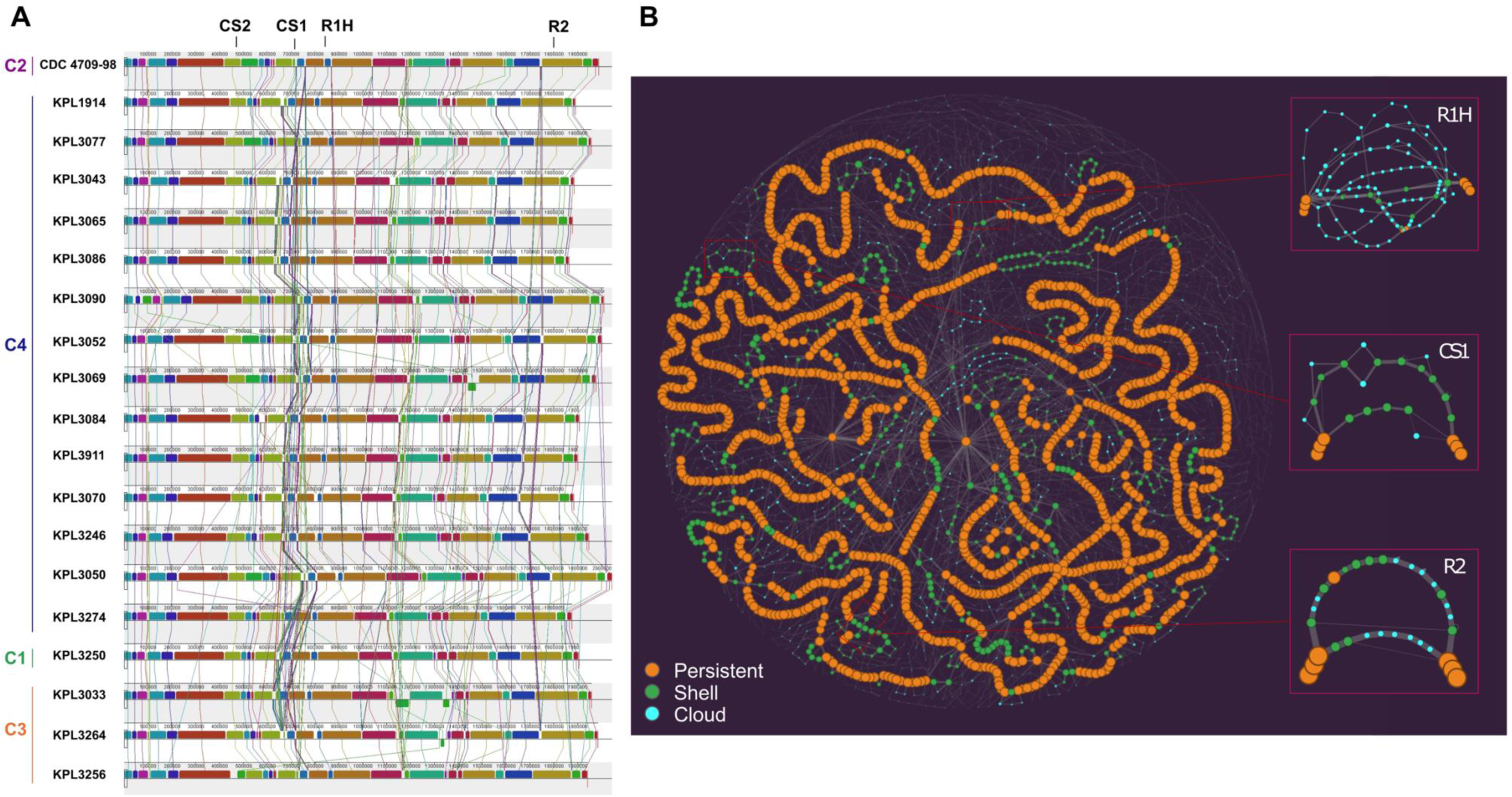
*D. pigrum* displays conserved chromosomal synteny. **(A)** A MAUVE alignment of 19 closed *D. pigrum* genomes, with representatives from the four major clades in Fig. 1, showed a conserved order of chromosomal blocks across the phylogeny of strains collected 20 years apart. Vertical bars represent clades: clade 1 green, clade 2 purple, clade 3 orange, and clade 4 blue. CS1 and CS2 designate the CRISPR-Cas sites (Fig. 8), R1H represents the RM system insertional hotspot and R2 represents the site containing either a Type II m5C RM system or a Type IV restriction system (Fig. 7). **(B)** This PPanGGOLiN partitioned pangenome graph displays the overall genomic diversity of the 28 *D. pigrum* genomes. Each graph node corresponds to a GC; node size is proportional to the total number of genes in a given cluster; and node color represents PPanGGOLiN assignment of GCs to the partitions: persistent (orange), shell (green) and cloud (blue). Edges connect nodes that are adjacent in the genomic context and their thickness is proportional to the number of genomes sharing that neighboring connection. The insets on the right depict subgraphs for sites R1H, CS1 and R2 showing several branches corresponding to multiple alternative shell and cloud paths. These sites with higher genomic diversity are surrounded by longer regions with conserved synteny, i.e., long stretches of consecutive persistent nodes (GCs). The static image depicted here was created with the Gephi software (https://gephi.org) (128) using the ForceAtlas2 algorithm (129) with the following parameters: Scaling = 20,000, Stronger Gravity = True, Gravity =6.0, LinLog mode = True, Edge Weight influence = 2.0.

### *D. pigrum* has a core genome that has leveled off, an open pangenome, and a high degree of conservation at the amino acid and nucleotide level

Analysis of all 28 *D. pigrum* genomes revealed a conservative core of 1,102 single-copy GCs, as defined by the intersection of results from three algorithms including bidirectional best hits (BDBH) **(File S1, Fig. Bi)**. A core of 1,134 GCs was defined by the intersection of two algorithms when BDBH was excluded **(File S1, Table A and Fig. Bii).** The *D. pigrum* core genome has leveled off in size (**Fig. 3A**). Meanwhile, the pangenome continued to increase with each additional genome reaching 3,700 GCs (**Fig. 3B**), of which 30.6% (1,134/3,700) are core (**File S1, Fig. Bii**). The average number of coding sequences (CDS) per genome was 1765 and, on average, the core constituted ∼64% (1,134/1,765) of the CDS in each individual genome (**File S1, Table A**). These results from GET_HOMOLOGUES (42) generally agreed with those from Anvi’o (52, 53), allowing us to leverage Anvi’o for additional analyses. In the Anvi’o-derived single-copy core (38.2%; **File S1, Fig. C**), 89.4% (993/1,111) of the GCs had a functional homogeneity index score equal to or higher than 0.98 indicating a high degree of conservation at the amino acid level. This fits with an average nucleotide identity (ANI) over 97.58% for all 28 genomes (**File S1, Fig. Biii**), matching earlier findings with 11 strain genomes (28). Moreover, two sets of three recently collected strains each shared over 99% ANI, as well as similar accessory Clusters of Orthologous Group (COG) annotations (**File S1, Fig. D**). This revealed highly similar strains in the nasal microbiota of different individuals in Massachusetts. Of these, two strains collected from different people were nearly identical (**Fig. 1**).

**Figure 3.**
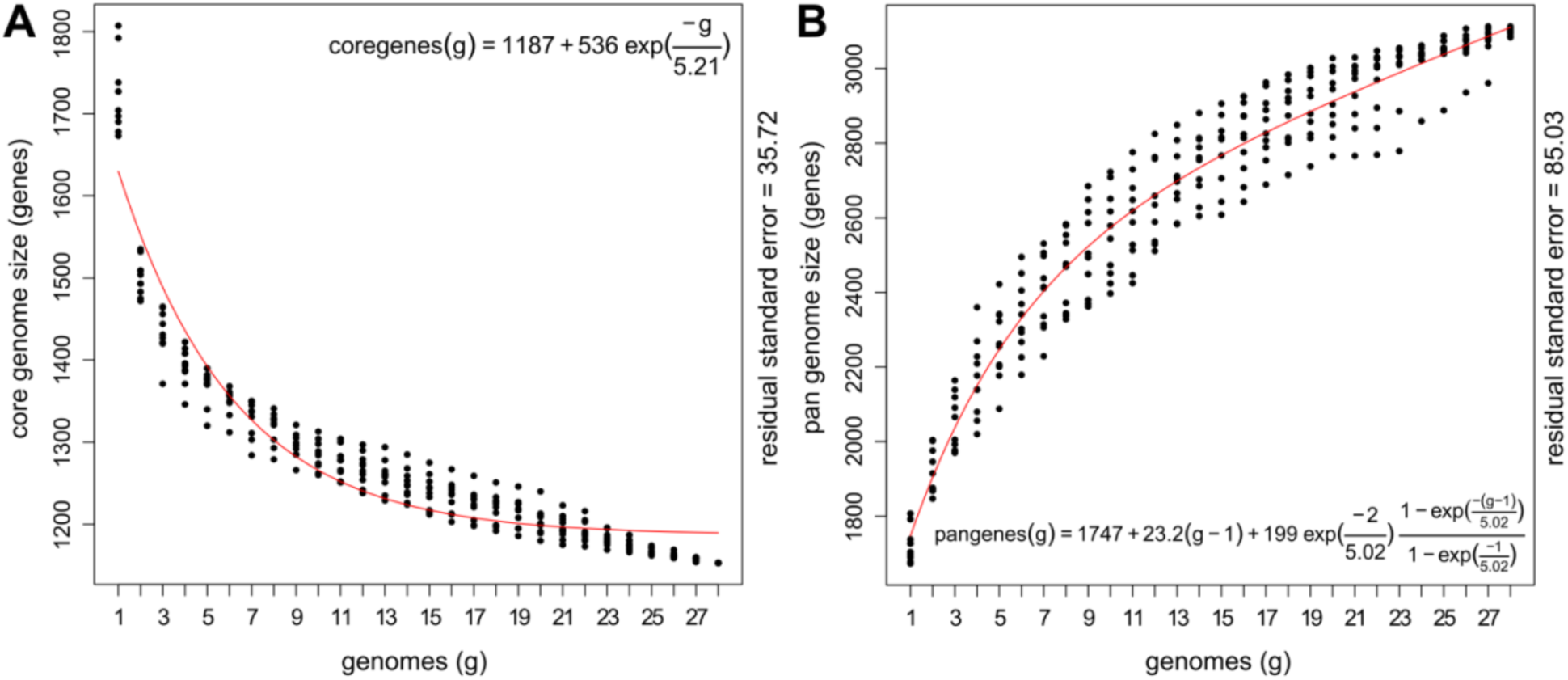
The *D. pigrum* core genome levels off and the pangenome remains open. **(A)** The *D. pigrum* core (n=28) genome started to level off after 17 genomes, as predicted using a Tettelin curve fit model (red line). Whereas, with 28 genomes, the **(B)** pangenome continued to increase in gene clusters with each additional genome. *D. pigrum* **(A)** core and **(B)** pangenome size estimations were based on ten random genome samplings (represented by black dots) using the OMCL algorithm defined gene clusters in GET_HOMOLOGUES v. 3.1.4.

### The *D. pigrum* accessory genome is enriched for gene clusters involved in mobilome and host defense

Of the 49,412 individual genes identified across the 28 genomes, 63.8% (31,501/49,412) had informative calls to a single functional COG annotation (i.e., their assignment corresponds to a single COG category other than S or R) (54, 55) (**Fig. 4A**). Using Anvi’o, we observed that GCs involved in mobilome, in defense mechanisms and in carbohydrate transport and metabolism were overrepresented in the accessory compared to the core genome (**Fig. 4B**). GCs classified to these three COG categories accounted for 3.9%, 6.6%, and 8.5% of the *D. pigrum* accessory genome, respectively. The proportion of accessory functions was similar among all strains, but the size of their accessory genomes varied (**File S1, Fig. D**). Because genome stability is relevant to suitability of a candidate beneficial microbe for therapeutic use, we focused subsequent analysis on the predicted mobilome and defense mechanisms.

**Figure 4.**
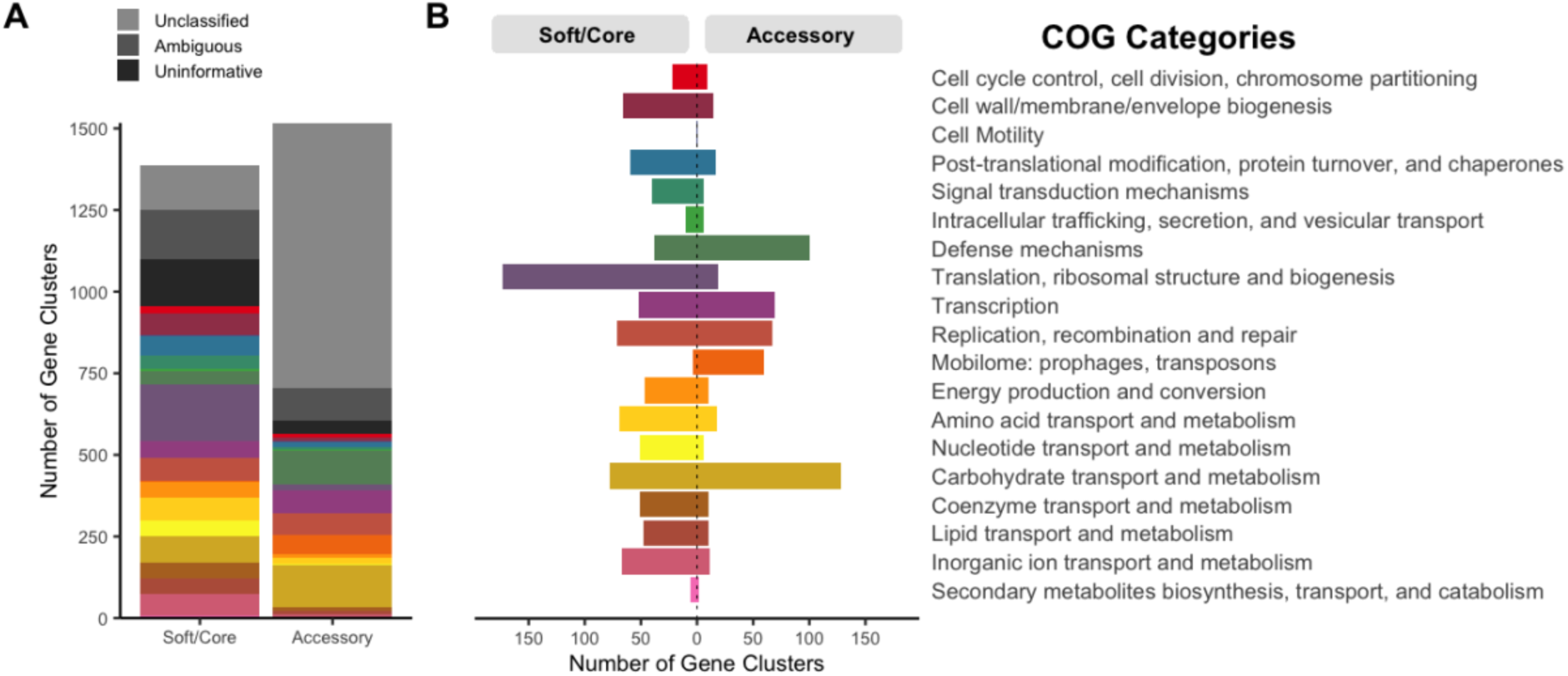
The accessory genome of *D. pigrum* has functional enrichment for defense mechanisms, mobilome, and carbohydrate transport and metabolism genes. (**A**) Out of the total 49,412 individual genes identified across the 28 analyzed genomes, up to 8,242 genes (16.7%) lacked a COG annotation, 5,221 (10.6%) had an ambiguous COG category annotation (more than one COG category), and 4,448 (9.0%) had an uninformative annotation (belong to the S or R COG category). At the gene cluster (GC) level, only 37.2% of the 1,517 GCs present in the accessory genome had an informative COG assignment compared to 68.7% of the 1,388 GCs in the soft/core. **(B**) The number of GCs present in the accessory genome was several folds higher than in the soft/core for the following informative COG assignments (colored categories): defense mechanisms (olive, 2.60 fold), mobilome: prophages, transposons (orange, 14.88 fold) and carbohydrate transport and metabolism (khaki, 1.66 fold). This was determined using the COG functional annotations defined in our Anvi’o analysis of the soft/core (“core” and “soft core” bins) versus accessory (“shell” and “cloud” bins). Since many GCs have individual genes with distinct COG annotations each individual gene was counted as 1/x with x being the number of genes in each GC.

### *D. pigrum* hosts distinct integrated phage elements, insertional elements and a group II intron

Of the total GCs in the pangenome, 2.2% were predicted to be part of the mobilome. MGEs can negatively affect genome stability and can positively affect strain diversification. Therefore, we systematically searched for various types of MGEs, including phage elements, plasmids and insertional elements that interact with *D. pigrum.* First, using the Phage Tool Enhanced Release (PHASTER) database (56, 57), we identified four distinct, and mostly intact, integrated phage elements, i.e., prophages (**Fig. 5**). We gave these the provisional names *Dolosigranulum* phage L1 through L4. All four were in the size range common for Firmicute phages and had a life-cycle-specific organization of its CDS with lytic and lysogenic genes separated (**Fig. 5**) (58–60). Predicted prophage L1 from *D. pigrum* KPL3069 was the most intact with two attachment (*attP*) sites and an intact integrase most similar to that of the *Streptococcus* prophage 315.2 (NC_004585; E-value 7.85e-69) (61). Prophages L2 and L3 from *D. pigrum* KPL3090 also had intact integrases, with similarity to other streptococcal phages, but lacked distinguishable *attP* sites. Beyond these similarities, other CDS from L1-4 displayed few and dissimilar matches to known phage elements (**File S2**) indicating that *D. pigrum* hosts a distinct set of lysogenic phage that are expected to have a limited host range.

**Figure 5.**
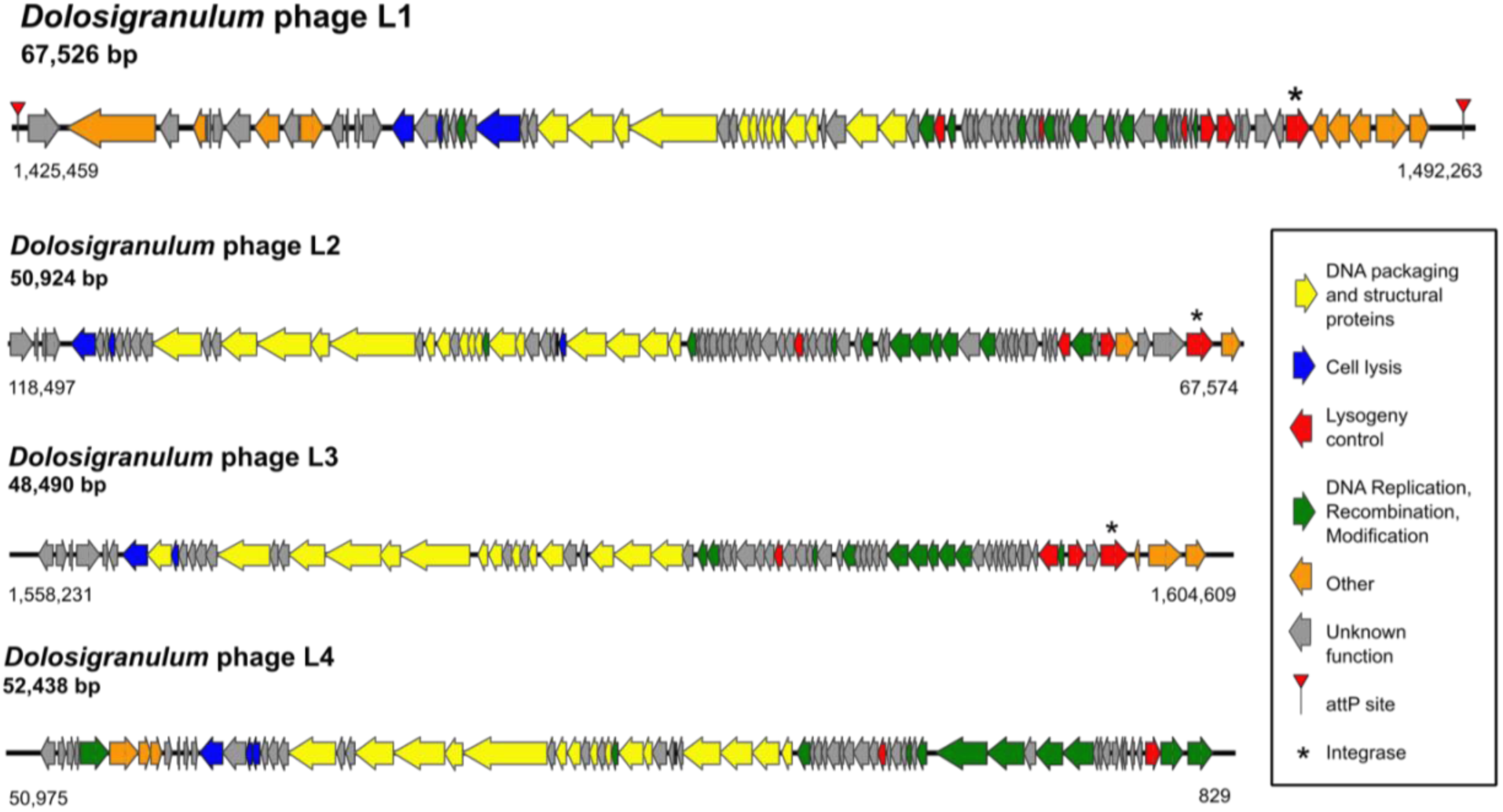
*D. pigrum* has an intact prophage. Map of the four predicted phages: *Dolosigranulum* phage L1 from KPL3069; L4 from KPL3256; L2 and L3 from KPL3090. The most complete phage was L1 from KPL3069 with an intact integrase and two *attP* sites. All the putative phages exhibited a typical life-cycle-specific organization with lytic genes on one side and lysogenic genes on the other. We detected phage elements using the PHASTER databased on 11/8/2018

Second, using the Gram-positive plasmid database PlasmidFinder (62), we detected no autonomous plasmids. However, a nearly complete fragment of the *S. aureus* plasmid pUB110 is integrated in the chromosome of four strains and includes a gene encoding for kanamycin resistance (**File S2, Fig. B**). This prompted a systematic search for antibiotic resistance genes using the Comprehensive Antibiotic Resistance Database in the Resistance Gene Identifier (CARD-RGI) (63, 64). Six of the twenty-eight genomes are predicted to encode antibiotic resistance genes for erythromycin and/or kanamycin, which are located within a CRISPR array or the integrated plasmid, respectively (**File S2**).

Third, we identified GCs predicted to be either transposases (eight) or integrases (five) using a multistep approach (**File S2, Table A**). Transposases are thought to function both as detrimental, selfish genetic elements that can disrupt important genes and as diversifying agents that can provide benefit to host cells through gene activation or rearrangements (65, 66). Among the 26 genomes containing at least one transposase CDS the mean was 4.42 (median 3.5) with a maximum of 13 per genome. Transposases were more prevalent and abundant than integrases (**File S2, Table A**). One of the predicted transposases was the GC containing the third largest number of sequences. This is consistent with reports that genes encoding transposases are the most prevalent protein-encoding genes detected across the tree of life when accounting for both ubiquity and abundance (67). We detected 74 intact instances of this most common transposase, an ISL3 family transposase with similarity to ISSau8, across 22 of the *D. pigrum* genomes with a mean (median) of 3.36 (2) and a maximum of 11 copies per genome (GC_00000003 **File S2, Table A**). As shown on the PPanGGOLin graph (**Fig. 6Ai**), this transposase is inserted at multiple different sites within and across the genomes (**Fig. 6Aii**; **File S2, Table A**). The most common of these is likely the ancestral insertion site (top, **Figure 6Bi**). The absence of a cotraveling CDS is consistent with this ISL3 family transposase being part of an insertional sequence (IS). Following standards for IS nomenclature, we propose the name ISDpi1 (66).

Fourth, the PPanGGOLin graph (68) revealed insertion of a predicted group II intron reverse transcriptase-maturase at multiple sites across multiple *D. pigrum* genomes (**Fig. 6Aiii**; **File S2, Table A**). Group II introns are MGEs commonly found in bacterial genomes that consist of a catalytic RNA and an intron-encoded protein that assists in splicing and mobility (69). Like transposases, group II introns can play both detrimental and beneficial roles within their host. We detected this intron-encoding GC in all 28 genomes with a mean (median) of 4.7 (3.5) and range of 1 to 14 copies per genome. This GC contained the highest number of individual gene sequences of any GC with 132 (GC_00000001 **File S2, Table A**). It is most closely related to the bacterial class C intron-encoded protein from La.re.I1 in *Lactobacillus reuteri* with 44% identity and 65% similarity over 419 amino acids (70). These data are consistent with an intact bacterial reverse transcriptase/maturase expected to facilitate splicing and mobility of the group II intron (69).

### A systematic search identifies multiples types of defense systems to protect *D. pigrum* from MGEs

The enrichment for defense mechanisms in the accessory genome of *D. pigrum* is combined with the relative paucity of plasmids and prophages among *D. pigrum* genomes. Based on this, we performed a systematic search of the pangenome for known bacterial host defense systems, including RM, deity-named defense and CRISPR-Cas systems.

### *D. pigrum* harbors a diverse collection of RM systems

In bacteria, individual RM systems can differ with respect to target sequence, active site architecture, and reaction mechanisms, but all recognize the methylation status of target sequences on incoming DNA and degrade inappropriately methylated (non-self) DNA. Type I-III systems largely recognize and digest a target sequence when it lacks the appropriate methyl group. In contrast, Type IV systems, which lack a methyltransferase, are composed of a methyl-dependent REase that cuts a target sequence when it contains a specific methyl-modification. RM systems and their recognition sequences are often strain specific. Therefore, we characterized and compared the repertoire of RM systems present in each of the 19 *D. pigrum* strains sequenced via SMRTseq, defining the methylome of each strain using SMRTseq kinetics (Basemod analysis) and predicting the recognition sequences of each system via REBASE analysis (71) (**Fig. 7A**; **Table S2**; **File S2**). Most of the modifications detected were m6A with only one m4C being found. There were several genes coding for m5C enzymes, but their products are not usually detected by the PacBio software. Only one positive m5C enzyme was identified. Among the RM systems, most were Type II, although half the strains had a Type IV enzyme of unknown specificity. The Type I-III systems were associated with 19 individual target recognition motifs identified by methylome analysis (**Fig. 7A; Table S2; File S2**).

**Figure 6.**
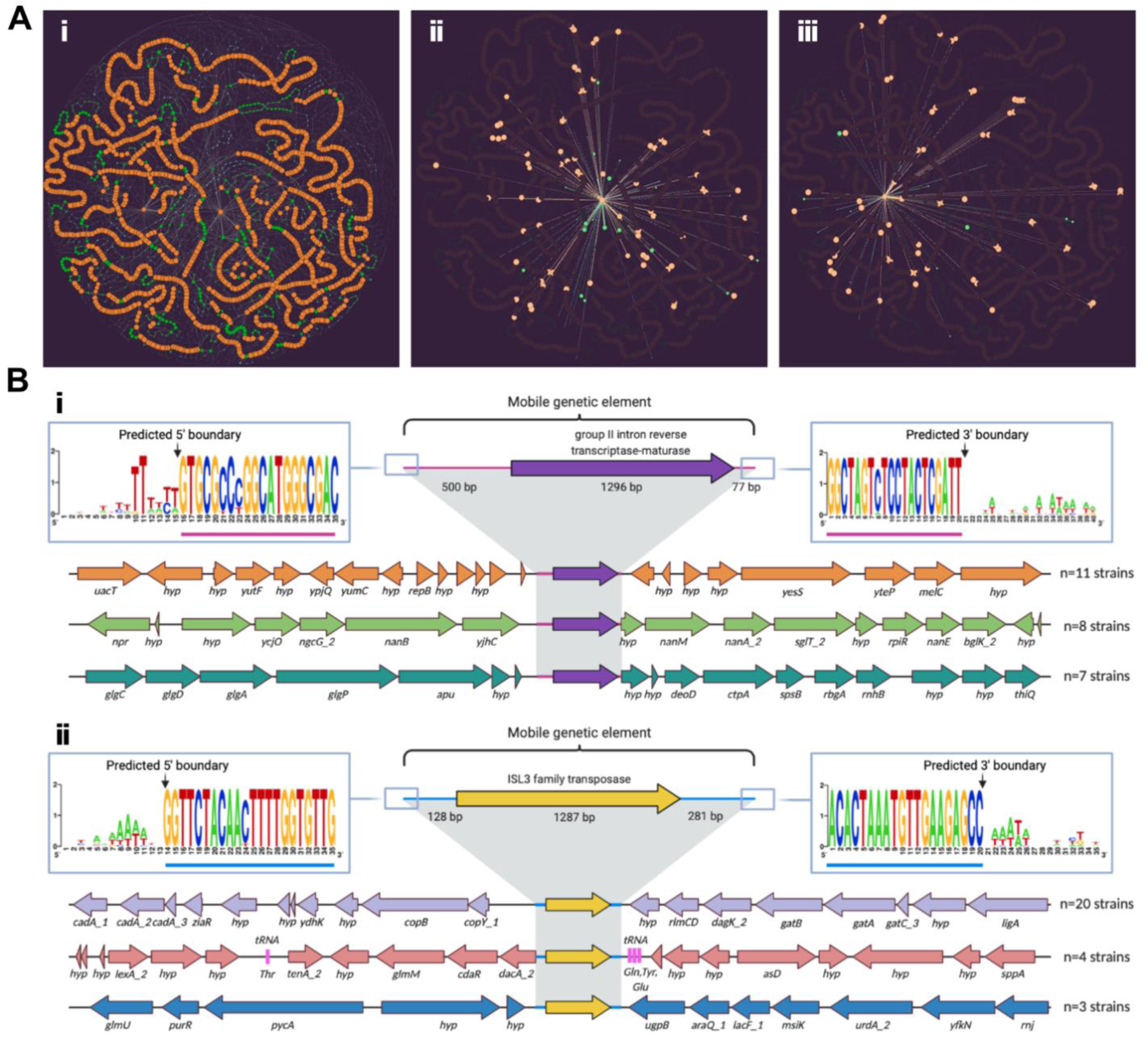
*D. pigrum* genomes host a few highly prevalent MGEs. (**Ai**) On the PPanGGOLiN partitioned pangenome graph for the 28 *D. pigrum* genomes, (**Aii**) we highlight the neighboring connections for the persistent GC of a predicted group II intron reverse transcriptase-maturase (purple in **Bi**) and (**Aiii**) a predicted ISL3 family transposase (yellow in **Bii**). Each graph node corresponds to a GC; node size is proportional to the total number of genes in a given cluster; and node color represents PPanGGOLiN assignment of GCs to the partitions: persistent (orange), shell (green) and cloud (blue). Edges connect nodes that are adjacent in the genomic context and their thickness is proportional to the number of genomes sharing that neighboring connection. In **Aii** and **Aiii**, only the adjacent neighboring nodes and edges for each of the depicted GCs are contrast colored against the background pangenome graph. (**B**) The most common genomic neighborhoods, respectively, for the predicted (**Bi**) group II intron reverse transcriptase-maturase and (**Bii**) ISL3 family transposase. BOFFO (https://github.com/FredHutch/boffo) identified the chromosomal coordinates of each MGE integration event in individual strains, and groupings of co-located genes residing within the same neighborhood structure across strains were visualized using Clinker (https://github.com/gamcil/clinker). ClustalOmega alignments of flanking regions across groupings revealed predicted terminal sequence boundaries (consistent 5’-3’ sequences across integration events) for each MGE. The three most common genomic loci for each MGE were rendered using BioRender.

**Figure 7.**
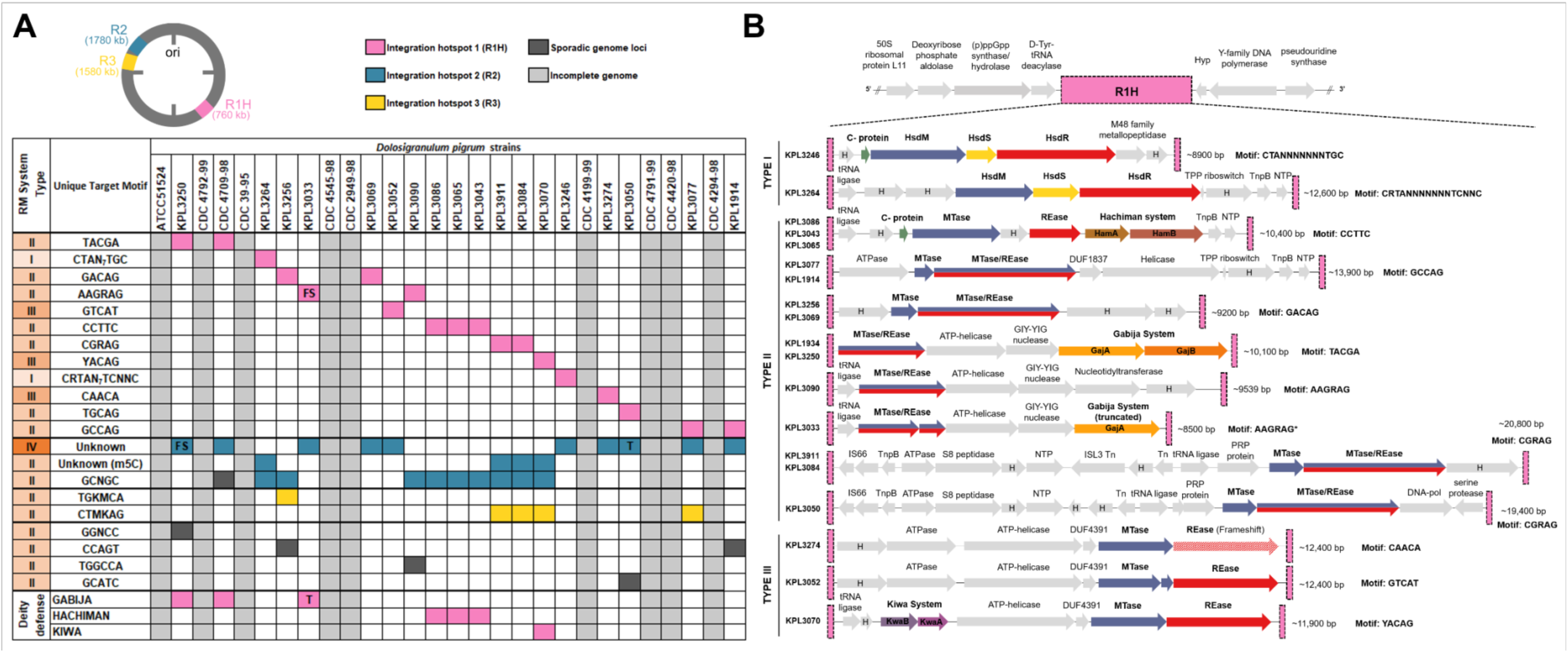
*D. pigrum* hosts a diverse collection of restriction modification (RM) systems at three distinct loci. (**A**) Conserved methyl-modifications associated with RM defense systems of *D. pigrum* strains. White and light grey cells indicate that a modified motif was not detected or no SMRTseq data was available for a specific strain, respectively. Colored cells indicate that a motif was detected and the approximate genomic loci of the RM system responsible across strains are indicated with pink (R1H), blue (R2) or yellow (R3) cells. Sporadic occurrences of RM systems that do not appear conserved in more than a single strain are indicated by dark grey cells. (**B**) The organization of gene clusters within RM system integrative hotspot 1 (R1H), which harbors a diverse collection of RM systems, including Type I (n=2), Type II (n=6) and Type III (n=3), in addition to other mobile elements/transposons systems, including Hachiman, Gabija, and Kiwa defenses. R1H is flanked upstream by region containing genes for (p)ppGpp synthase/hydrolase and D-Tyr-tRNA (Tyr) deacylase proteins, and downstream by a region with genes for Y-family DNA polymerase and an rRNA pseudouridine synthase protein. Hypothetical genes are indicated by grey arrows assigned ‘H’.

### The *D. pigrum* Type IV RM system is inversely related to a specific m5C-associated Type II system

We noted an inverse relationship between the presence of the *D. pigrum* Type IV REase and a specific m5C-associated Type II RM system that modified the second cytosine residue within the motif GCNGC (**Fig. 7A**). This inverse relationship was found to be interdependent between strains based upon a Fisher exact test (*p*=0.0055). The Type II m5C system was present in nine *D. pigrum* genomes that lacked the Type IV REase. Conversely, the Type II m5C system was absent in eight strains that contained the Type IV REase. Type IV REase that target m5C-modified motifs have the potential to limit the spread of RM systems that utilize m5C modifications. The *D. pigrum* Type IV REase appears related (99%coverage / 43% identity) to *S. aureus* SauUSI, a modified cytosine restriction system targeting S^5m^CNGS (either ^m5^C or ^5hm^C) where S is C or G. Based on the inverse relationship of the Type IV and Type II m5C systems, this strongly indicates that the *D.pigrum* Type IV system targets m5C containing sequences, including GCNGC, GGNCC and potentially the recognition sequence of the other m5C enzyme, M.Dpi3264ORF6935P.

### The Type IV and specific m5C-associated Type II RM systems are present at the same integration site

To decipher the basis for the inverse relationship between these two RM systems, we asked where each was incorporated in the *D. pigrum* genomes. In 18 of the 19 strains, the Type IV REase or the m5C-associated Type II system are inserted into the same genetic locus, dubbed R2 (**Fig. 2**; **File S2, Fig. Ci**). In the one strain that carried both the Type IV REase and the m5C-associated Type II system, CDC 4709-98, the Type IV is present at R2, whereas the m5C-system is integrated at an unrelated locus downstream from a tRNA-Leu site. MGEs that carry similar integrases tend to integrate at the same sites in the chromosome, but in most strains, we did not observe any integrase or additional genes co-occurring with the RM systems at this site.

### Many *D. pigrum* RM systems compete for an integration hotspot

Extending our analysis, we identified a genomic locus with an unexpectedly high frequency of variable genes across all 28 genomes. We dubbed this site RM system integration hotspot 1 (R1H), because it harbors a diverse collection with 12 different RM systems spanning Types I, II, and III across strains (**Fig. 2; Fig. 7B**). Co-occurring with these RM systems in R1H, we also identified three of the antiphage deity-named defense systems: Hachiman, Gabja, and Kiwa present across seven strains (**Fig. 7B**). A third RM system integration site (R3) contained two different Type II systems along with an IS66 transposase family of genes (**File S2, Fig. Cii**), consistent with the known association of defense systems and MGEs (72).

### *D. pigrum* encodes subtype II-A and I-E CRISPR-Cas systems

CRISPR-Cas systems provide adaptive/acquired defense (immunity) against MGEs (46). All of the complete *D. pigrum* genomes encoded at least one subtype II-A or I-E CRISPR-Cas system (**Fig. 8A**; **Table S3A**), based on the CRISPRDetect database (73). Of the 32 CRISPR-Cas systems detected, 22 are subtype II-A, which is mostly found in Firmicutes (74) and is the predominant CRISPR-Cas system among *Lactobacillus* (75). Subtype II-A (circles, **Fig. 8A**) and I-E (stars, **Fig. 8A**) CRISPR-Cas systems were generally intermixed within the four major clades, although two distal clades harbored only one type. A single genomic locus (CS1) contained either a subtype II-A or a subtype I-E CRISPR-Cas system in all 19 closed genomes (**Fig. 2B**; **Figs. 8A-B**). A second CRISPR-Cas system (triangles, **Fig. 8A**) was found at a second location (CS2) in 4 of these 19 genomes, from three of the four clades (**Fig. 8B**).

**Figure 8.**
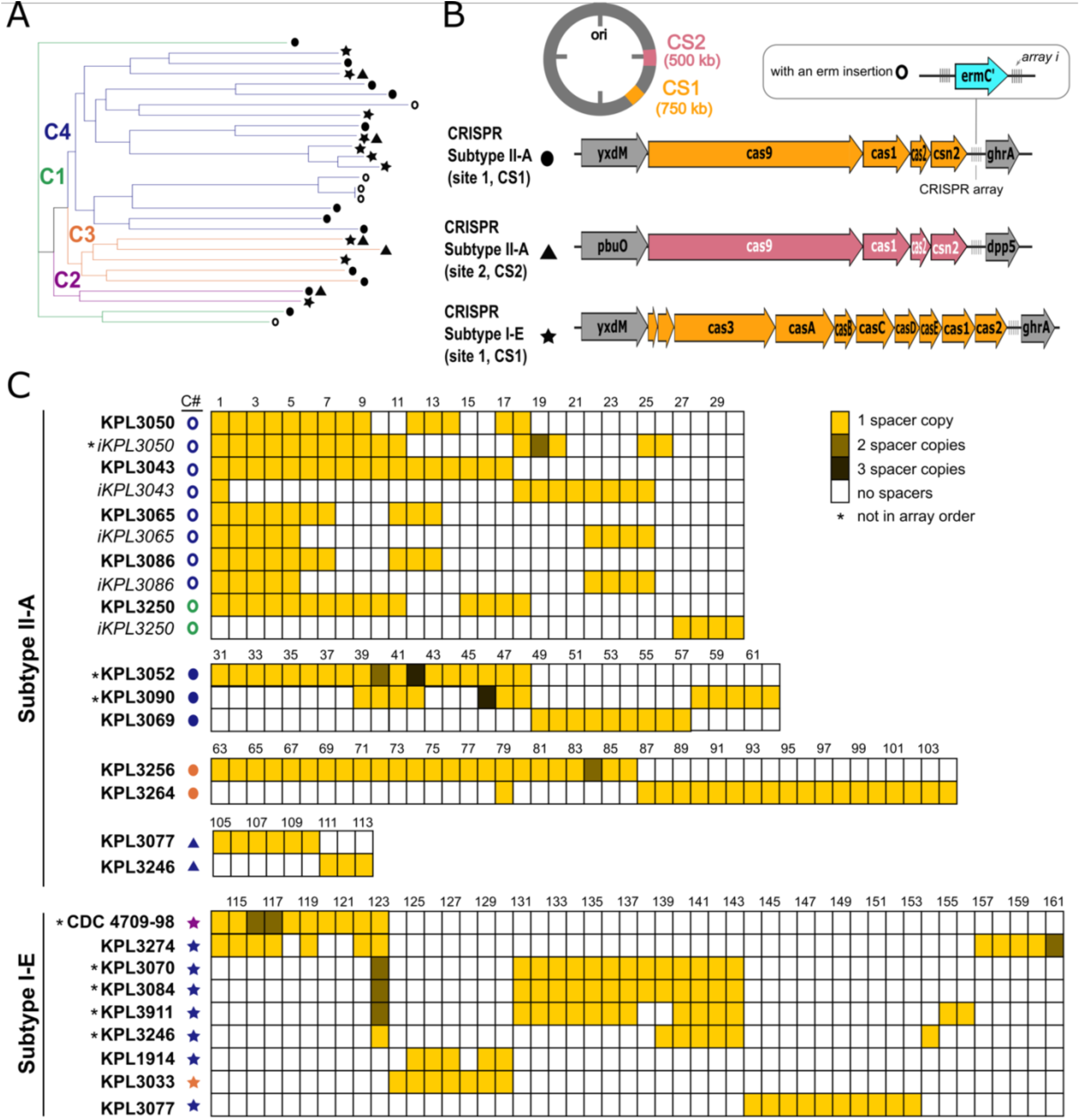
*D. pigrum* encodes subtype I-E and II-A CRISPR-Cas systems with a large but sparsely shared history of MGE invasion. (**A**) CRISPR-Cas subtype II-A (circles and triangles) and I-E systems (stars) were intermixed among strains in all four clades, with type II-A being most common (**Table S3A**). Two distal clades had only a subtype II- A system (KPL3043, KPL3065-KPL3086, KPL3090, KPL3052 and KPL3069) or a subtype I-E system (KPL3070, KPL3084 and KPL391). Three genomes (KPL3077, KPL3246 and CDC 2949-98) have both types of system, with each at a different locus. **(B)** The most common location, CRISPR-Cas system insertion site (CS1), is between the ABC transporter permease protein (*yxdM*) and the glyxyolytate/hydroxypyruvate reductase A (*ghrA*) genes. However, subtype II-A systems are also found in between the guanine/hypoxanthine permease (*pbuO;* NCS2 family permease) and dipeptidyl-peptidase 5 (*dpp5;* S9 family peptidase) genes at CRISPR-Cas insertion site 2 (CS2). Five of the strains with a subtype II-A system in CS1 had a predicted rRNA adenine N-6- methyltransferase (*ermC*’) gene integrated in their CRISPR arrays (open circles) (**C**) Representation of the spacers (**Table S3B; File S2**) found among the different CRIPSR systems in the 19 closed genomes. We found 161 unique spacers, less than one third of which were homologous to phages and plasmids found among other Firmicutes. Strains KPL3050, KPL3250, KP3086-KPL3065, and KPL3043 shared the most spacers among the subtype II-A CRISPR-Cas system, with the distal clade of with KPL3043 and KPL3065-KPL3086 sharing 15 spacers. The distal clade with KPL3070, 3084, and 3911 shared the most spacers (12) among the subtype I-E system. CRISPR-Cas systems and spacers hits were determined using the CRISPRdetect and CRISPRtarget database on 2/16/2019 while shared spacers were determined using CRIPSRCompar on 3/18/2019.

### *D. pigrum* CRISPR-Cas spacers point to undiscovered *D. pigrum* MGEs

Each of the 19 closed genomes included at least one complete CRISPR array. (As expected, most of the arrays were incomplete in the unclosed genomes.) Examining the CRISPR arrays in the 19 closed genomes revealed two key findings. First, the spacer sequences predict the existence of a diversity of undiscovered *D. pigrum* phages and plasmids with a mean (median) number of spacers per array of 13 (12.5) for subtype II-A and 11.1 (12) for subtype I-E (**Table S3A**). Second, spacer sequences show a sparsely shared history of exposure to many MGEs (**Fig. 8C**; **Table S3A-B**). Only 60 of the 161 unique identified spacers were shared by more than one strain (**Fig. 8C**). The exceptions to this limited shared history were two distal clades with shorter branch lengths within Clade 4, which shared 15 and 12 spacers, respectively. Of these 27 spacers, 9 had similarity to known MGEs (**File S2**). A few other shared spacers were scattered among *D. pigrum* strains outside of these two distal clades. For example, *D. pigrum* KPL3033 (Clade 3) and KPL1914 (Clade 4) shared five spacers (**Fig. 8C**), one of which matched to the *Clostridium* phiCDHM19 phage (LK985322; spacer 129) (76). These shared spacers suggest strains within the host-range of specific MGEs. Spacer similarity to known MGEs indicated prior *D. pigrum* exposure to phage and plasmid elements that might be related to those found in other genera of Firmicutes, e.g., *Clostridium*, *Lactococcus*, *Streptococcus*, *Staphylococcus*, and *Enterococcus*. However, only 46/161 spacers had significant matches (match score >15) to previously identified MGEs, indicating that that *D. pigrum* CRISPR-Cas systems likely target a variety of yet-to-be-identified host-specific *D. pigrum* plasmids and phages, such as the predicted prophages in Figure 5.

## DISCUSSION

Multiple recent studies of the composition of human nasal microbiota identify *D. pigrum* as a candidate beneficial bacterium (1–30). Our systematic analysis of 28 *D. pigrum* strain genomes, including 19 complete and closed genomes, reveals a phylogeny in which strains collected 20 years apart intermingled in clades and showed remarkable stability in genome structure (**Figs. 1 and 2**). We confirmed that both the nucleotide sequence (≥97.5%) and chromosomal structure, aka synteny (**Fig. 2**), of the core genes are very highly conserved in *D. pigrum* (28). However, *D. pigrum* also has an open pangenome (**Fig. 3B**) with strain-level variation driven by gene gain/loss in variable regions located between the large blocks of syntenic core genes (**Fig. 2**). The *D. pigrum* accessory genome is enriched for GCs that include those involved in mobilome and defense mechanisms (**Fig. 4B**). A systematic search for MGEs identified no autonomous plasmids, a few prophages, and a small number of transposases and integrases. Among these are the first predicted intact *D. pigrum* prophage (**Fig. 5**), IS and group II intron (**Fig. 6**). In contrast, a systematic search for gene clusters involved in defense systems identified a diverse collection of RM systems, several deity-named defense systems (**Fig. 7**) and two types of CRISPR-Cas systems (**Fig. 8**) inserted at conserved sites across the genomes. Analysis of CRISPR spacer sequences points to a variety of previously unknown *D. pigrum* MGEs against which *D. pigrum* appears to effectively defend itself, which likely contributes to its overall genomic stability.

Many of the older *D. pigrum* strains were collected in the context of human disease (48) making it unclear whether these strains were contributors to disease, bystanders, or contaminants. In a previous analysis, we detected no virulence factors in the genomes of nine of these older strains from Laclaire and Facklam (48), consistent with *D. pigrum* having a commensal or mutualistic relationship with humans (28). Adding further support for this, we detected no virulence factors in any of our newer strains here (**File S2**), which were all isolated from healthy volunteers. Plus, many of the older strains are closely related in the phylogeny with these recent healthy-donor-derived strains (**Fig. 1**). These findings are consistent with there being only a few isolated reports of *D. pigrum* growth in samples from different types of infections (77–82). Of these, the repeated detection of *D. pigrum* alone in keratitis/keratoconjunctivitis raises the possibility that some strains might be rare causes of eye surface infection (83–86). We recommend future genome sequencing of ophthalmic infection isolates to ascertain whether and how these vary from currently sequenced avirulent strains.

Our results show that strain-level variation in *D. pigrum* is driven by gene gain/loss in variable regions located between large blocks of syntenic DNA (**Fig. 2**). This pattern is consistent with Oliveira and colleagues’ findings for the chromosomal structures of 80 different bacterial species (87). Furthermore, 19 closed genomes show the order of syntenic blocks of core genes in *D. pigrum* is conserved (**Fig. 2A**). *D. pigrum* has an average genome size of 1.93 Mb (median 1.91 Mb) (**File S1, Table A**) with an open pangenome (**Fig. 3B**). About 64% of each *D. pigrum* strain genome consists of core CDS, whereas, only about 30% of the *D. pigrum* pangenome consists of core GCs. This is similar to the percentage of core genes in the pangenomes of other colonizers of the human upper respiratory tract, such as *Staphylococcus aureus* (36%) and *Streptococcus pyogenes* (37%) (88).

HGT, much of it likely mediated by MGEs, plays an important role in strain diversification in free-living bacteria. However, a systematic search identified few such elements per genome among *D. pigrum* strains. In terms of MGEs that commonly mediate HGT, we detected no autonomous plasmids. However, we identified one complete and three partial predicted prophages (**Fig. 5**) among 27 distinct strain genomes (2 of the 28 genomes were almost identical). To our knowledge, the predicted complete prophage (L1) is the first phage element identified in *D. pigrum*. The disparate nature of these candidate prophages compared to those in current databases is consistent with *D. pigrum* having its own specific pool of yet-to-be-identified phage predators, consistent with the strain-level specificity of many known phages. This is further supported by the scarce homology of the phage spacers in the CRISPR arrays to those available in the databases. However, some *D. pigrum* prophages might share a distant common ancestor with streptococcal phages, as almost one fifth L1’s and at least one third of L2’s (77/202) and L3’s (51/187) predicted genes shared the most similarities to *Streptococcus* phage genes (**File S2**). Based on our findings, we predict that phage elements targeting *D. pigrum* have a narrow host range, consistent with patterns exhibited by other Firmicutes-targeting phages, such as those targeting *Listeria* and *Clostridium difficile* (58, 76).

In terms of MGEs that commonly move within genomes, *D. pigrum* genomes host a group II intron and most also host a small number of predicted transposase and/or integrases (**FileS2, Table A**). Once present in a genome, IS movement can lead to phenotypic variation among closely related strains through disruption of ORFs or changes in transcript due to insertion in or adjacent to promoters (65).

The small number of MGEs identified might be related to the multiple defense mechanisms present in each *D. pigrum* genome. RM systems are ubiquitous in bacteria and present in ∼90% of genomes (71). They play a key role in protecting bacterial genomes from HGT, including MGEs, and maintaining genome stability. The variety of RM systems within and among *D. pigrum* genomes is consistent with this role. To our knowledge, this is the first report of a strongly inverse relationship between an m5C-targeting Type IV REase and an m5C-associated Type II system within the same chromosomal locus. A similar relationship was described previously for two antagonistic Type II systems in *Streptococcus pneumonia*, where strains possess either DpnI (which cleaves only modified G^m6^ATC) or DpnII (which cleaves only unmodified GATC) (89). It remains unclear whether the inverse relationship observed between the two *D. pigrum* systems results from competition for an integration hotspot within a *D. pigrum* genome (R2; **Fig. 7**) or whether the Type II systems m5C-modified target motif is incompatible with the Type IV REase. Determination of the exact underlying mechanism for this TypeIV/TypeII relationship warrants future investigation and has implications for other bacterial genomes.

CRISPR-Cas systems are another common bacterial defense system that maintain genomic stability. In a recent analysis of complete genomes from 4010 bacterial species in NCBI RefSeq, 39% encode *cas* clusters (74). Several characteristics of the predicted *D. pigrum* CRISPR-Cas systems suggest these are active. First, the preservation of repeats and spacers along with all of the core Cas gene suggests active systems, since inactive systems often show evidence of degeneration in terms of inconsistent repeat/spacer lengths (75). Second, the diversity of spacers among *D. pigrum* strains supports the likelihood of activity (90). *D. pigrum* belongs to the order *Lactobacillales* in the phylum Firmicutes. Similar to our observations in *D. pigrum*, among 171 *Lactobacillus* species, when multiple CRISPR-Cas systems are present in a single genome these are most often a subtype I-E and subtype II-A, and these two subtypes predominate among Type I and II systems in *Lactobacillus* (75). More broadly, there is a positive association between subtype I-E and subtype II-A systems within the phylum Firmicutes (74). Within *Lactobacillus*, type I systems contain the longest arrays (average 27 spacers) (75) and we see something similar among the *D. pigrum* strains. Of the spacers with matches to known plasmid and phage elements in the Genbank-Phage, Refseq-Plasmid, and IMGVR databases in CRISPRTarget, almost half of the identified spacers corresponded to plasmid elements. Subtype II-A systems in *Lactobacillus* actively transcribe and encode spacers that provide resistance against plasmid uptake based on plasmid interference assays in which an exogenous plasmid is engineered to contain endogenous spacer sequences (75, 91). This defense mechanism might explain the lack of autonomous plasmids in *D. pigrum* strain genomes to date.

The majority of *D. pigrum* CRISPR spacers lack homology to known MGEs. This is consistent with a large-scale analysis of bacterial and archaeal genomes in which only 1 to 19% of spacers (global average ∼7%) in genomes match known MGEs, mostly phages and plasmids and uncommonly to self. Also, spacers without a match share basic sequence properties with MGE-matching spacers pointing to species-specific MGEs as the source for CRISPR spacers (92). In this context, our findings indicate *D. pigrum* strains defend themselves against a wealth of yet-to-be-identified *D. pigrum-*specific MGEs. Some of these MGEs might be key to developing a system for genetic engineering of *D. pigrum*.

Like other pangenomic studies, this one has both general and species-specific limitations. First, the open pangenome indicates that the accessory gene space of *D. pigrum* remains to be more completely assessed through sequencing strains beyond the 28 investigated here. All but 1 of these 28 strains were collected in North America, so a next step is genome sequencing *D. pigrum* isolates from human volunteers from diverse geographic settings on other continents. Second, many more isolates would need to be collected over time to generate a comprehensive analysis of *D. pigrum* strain circulation in humans across the U.S., and beyond. Third, this is a systematic computational prediction of genome defense systems and MGEs. The next step is experimental verification of the function of these computationally predicted entities, which underscores the need for a system to genetically engineer *D. pigrum*. Fourth, here, we systematically identified known genomic elements that can affect bacterial genomic stability. This leaves a large proportion of *D. pigrum’s* accessory genome to be explored in future work.

In conclusion, a growing number of studies point to *D. pigrum* as a candidate beneficial bacterium with the potential for future therapeutic use to manage the composition of human nasal microbiota to prevent disease and promote health (40). One standard for bacterial strains for use in humans, either in foods, the food chain or therapeutics, is the absence of antimicrobial resistance (AMR) genes against clinically useful antibiotics (93). A prior report of 27 *D. pigrum* strains shows all are susceptible to clinically used antibiotics with the exception of frequent resistance to erythromycin (48). Consistent with this, only 6 of the 17 new *D. pigrum* genomes reported here encode AMR genes with predicted resistance to erythromycin and/or kanamycin (**File S2**). This confirms the broad antimicrobial susceptibility of *D. pigrum*. Further supporting its safety, we detected no virulence factors in these 28 genomes. Moreover, this pangenomic analysis of 28 *D. pigrum* isolates collected over the span of 20 years revealed remarkable stability in both strain circulation and chromosomal structure. Consistent with this stability, we detected relatively few MGEs in each genome; however, each genome hosted a variety of defense systems for protection against MGEs, and HGT in general. The antibiotic susceptibility, genomic stability, capacity for defense against HGT and lack of known virulence factors described here all support the safety of *D. pigrum* as a candidate for use in clinical trials to determine its potential for therapeutic use.

## MATERIAL AND METHODS

### Collection of new *D. pigrum* strains

We collected strains of *D. pigrum* from children and adults using supervised self-sampling of the nostrils with sterile swabs at scientific outreach events in Massachusetts in April 2017 and April 2018 under a protocol approved by the Forsyth Institutional Review Board (FIRB #17-02). All adults provided informed consent. A parent/guardian provided informed consent for children (<18 years old) and all children ≥5 years provided assent. (Self-sampling by children was considered evidence of assent.) Briefly, participants rubbed a sterile rayon swab (BBL; Franklin Lakes, NJ, USA) around the surface of one nasal vestibule (nostril) for 20 seconds, then immediately inoculated this onto BBL Columbia Colistin-Nalidixic acid agar with 5% sheep’s blood (CNA blood agar). After 48 hours of incubation at 37°C in a 5% CO_2_ enriched atmosphere, each CNA blood agar plate was examined and colonies with a morphology typical for *D. pigrum* were selected for purification. Purified isolates were verified to be *D. pigrum* by 16S rRNA gene colony PCR (GoTaq Green, Promega; Madison, WI, USA) using primers 27F and 1492R and Sanger sequencing from primer 27F (Macrogen USA; Cambridge, MA, USA).

### Genomic DNA extraction

All *D. pigrum* strains were cultured from frozen stocks on CNA blood agar plates at 37°C with 5% CO_2_ for 48 hours. For each strain, cells from eight plates were harvested with a sterile cotton swab (Puritan; Guilford, ME, USA) and resuspended in 1 ml of sterile 1X Phosphate Buffered Saline (PBS, Fisher; Waltham, MA, USA). Ten total 100 µl resuspensions were spread and grown on 47 mm, 0.22 µm-pore size polycarbonate membranes (EMD Millipore; Burlington, MA, USA) atop CNA agar plates at 37°C with 5% CO_2_ for 24 hours. Three membranes were resuspended in 20 ml of TES buffer (20 mM Tris-HCl Buffer, 1M, pH 8.0; 50mM EDTA; filter sterilized) and normalized to an OD_600_ of 1.0 +/-0.02. Half the resuspension was spun down at 5,000 rpm (2935 x g) for 10 min at 4°C. The genomic DNA was extracted using the Lucigen (Epicentre; Middleton, WI, USA) Masterpure Gram Positive DNA Purification Kit per manufacturer’s instruction with the following modifications: we increased the amount of Ready-Lyse lysozyme added per prep to 2.5 µl and deleted the bead beating step. The extracted genomic DNA was assessed for quantity using a Qubit per manufacturer instructions; for quality on a 0.5% agarose gel; and for purity by measuring 260/280 and 260/230 ratios on a Nanodrop spectrophotometer.

### Whole genome sequencing, assembly, and annotation

Single molecule, real-time sequencing (SMRTseq) was carried out on a PacBio Sequel Instrument (Pacific Biosciences; Menlo Park, CA, USA) with V2.1 chemistry, following standard SMRTbell template preparation protocols for base modification detection (www.pacb.com). Genomic DNA (5-10 µg) samples were sheared to an average size of 20 kbp via G-tube (Covaris; Woburn, MA, USA), end repaired and ligated to hairpin barcoded adapters prior to sequencing. Sequencing reads were processed using Pacific Biosciences’ SMRTlink pipeline (https://smrtflow.readthedocs.io/en/latest/smrtlink_system_high_level_arch.html) with the HGAP version 4.0 assembly tool standard protocol. Single contigs generated through HGAP were also processed through Circlator version 1.5.5 using default settings to assign the start site of each sequence to *dnaA* (94). All genomes were annotated with the NCBI’s Prokaryotic Genome Annotation Pipeline (PGAP) (95, 96) and uploaded to NCBI (accession: CP040408 - CP040424).

### Determination of the conservative core genome and the pangenome sizes

All the genomes were annotated with Prokka version 1.13.0 (97) prior to identification of the conservative core genome with GET_HOMOLOGUES version 3.1.4. (42, 98) using the cluster intersection (′compare_clusters.pl′, blastp) result of three algorithms: Bidirectional best-hits (BDBH), cluster of orthologs (COG) triangles (99), and Markov Cluster Algorithm OrthoMCL (OMCL) (100). The nucleotide level clustering for each of these algorithms was calculated with the ′get_homologues.pl′ script and the following parameters: -a ′CDS′, -A, -t 28, -c, -R, and either -G for COG, -M for OMCL, and no flag for BDBH. To obtain the nucleotide instead of the protein outputs, blastn instead of blastp was used to report clusters (parameter –a ′CDS′).

The pangenome was established using the OMCL and COG triangle algorithm with –t 0 parameter to get all possible clusters when running get_homologues.pl. The total clusters from the OMCL and COG pangenomes were then used by compare_clusters.pl with the –m flag to create a pangenome matrix tab file. The cloud, shell, soft core, and core genome of the isolates were then determined using the parse_pangenome_matrix.pl script in GET_HOMOLOGUES using the -s flag and the pangenome matrix tab file. The average nucleotide identity and genome composition analysis were also implemented (using the –A and –c parameters, respectively, in get_homologues.pl). For the genome composition analysis, which shows how many new CDS are added to the pangenome per new genome addition, a random seed (–R) of 1234 was selected.

### Phylogenomic tree construction

A core gene alignment was created for phylogenetic analysis using the nucleotide sequences from the conservative single-copy core GCs (n=1,102) identified with GET_HOMOLOGUES. These GCs were aligned with MAFFT version 7.245 (101) using default settings, renamed to match the isolate’s strain name, and concatenated into an MSA file through the catfasta2phyml.pl script using the concatenate (--concatenate) and fasta (-f) parameters (copyright 2010-2018 Johan Nylander). The core gene multiple sequence alignment was converted into a phylip file format with Seaview version 4.7 (102). An unrooted phylogenetic tree of the conservative single-copy core (**Fig. 1**) was generated using this phylip file and IQ-Tree version 1.6.9. (103). The ModelFinder function in IQ-Tree identified the GTR+F+ as the appropriate substitution model for tree construction (BIC value 5597954.8128) (104). Using this model, 553 maximum likelihood searches with 1000 ultrafast rapid Bootstraps (105) were used to generate the final maximum likelihood tree (ML= -2854949.911). A clade was defined as a monophyletic group of strains sharing a well-supported ancestral node.

### Synteny Analysis

We performed a whole genome sequence alignment on all closed genomes using progressive Mauve in Mauve version 2.4.0.r4736 with its default settings (50, 51). For the five genomes that we were unable to circularize, we manually fixed the start site to *dnaA* and added NNNNNNNNN to the region concatenating the ends of the contigs to mark it as a region of uncertainty in the synteny alignment. Manual curating was done with SnapGene version 4.2.11 GUI platform (SnapGene® software from GSL Biotech, <u>snapgene.com</u>).

### Functional analysis of the pangenome using Anvi’o

All genomes were re-annotated with an updated Prokka version (1.14.6) (97) with default parameters, including gene recognition and translation initiation site identification with Prodigal (106). The pangenome was analyzed using Anvi’o version 7 (52, 53). We followed the pangenome workflow to import Prokka annotated genomes into Anvi’o (http://merenlab.org/2017/05/18/working-with-prokka/), followed by the addition of functional COG annotations using the anvi-run-ncbi-cogs program with the --sensitive flag (runs sensitive version of DIAMOND (107)) and the 2020 updated COG20 database (108, 109). KEGG/KOfam (110, 111) and Pfam (112) annotations were also added to each genome .db file, as well as hmm-hits (113). The pangenome was calculated with the anvi-pan-genome program (flags: --minbit 0.5, --mcl-inflation 10 and --use-ncbi-blast) using blastp search (114), muscle alignment (115), ‘minbit heuristic’ (116) to filter weak hits, and the MCL algorithm (117). The functional and geometric homogeneity index, and the rest of the information shown in **File S1, Fig. C** were calculated following the standard Anvi’o pangenomic pipeline (http://merenlab.org/2016/11/08/pangenomics-v2). The core (n = 28), soft core (28 > n ≥ 26), shell (26 > n ≥ 3), and cloud (n ≤ 2) annotations from GET_HOMOLOGUES were added to the Anvi’o pangenomic database using the interactive interface. We defined the accessory as GCs present in ≤ 25 genomes and core as GCs present in ≥ 26 genomes. The output of this Anvi’o pangenomic analysis and the code used to generate it are available at https://github.com/KLemonLab/DpiMGE_Manuscript/blob/master/SupplementalMethods_Anvio.md. We used the summary file we exported from the Anvi’o pangenomic analysis to generate the functional enrichment plots shown in **Fig. 4** and **File S1, Fig. D** using an in-house R script (https://github.com/KLemonLab/DpiMGE_Manuscript/blob/master/SupplementalMethods_COGs.md) to wrangle and extract information on the informative COG20 annotated gene clusters (118, 119).

### <u>P</u>artitioned <u>PanG</u>enome <u>G</u>raph <u>O</u>f <u>Li</u>nked <u>N</u>eighbors (PPanGGOLiN) analysis

Gene clustering and annotation data were exported from the Anvi’o output and imported into PPanGGOLiN version 1.1.141 (68) to create a Partitioned Pangenome Graph (PPG) that assigned GCs to the ’persistent’, ’shell’, and ’cloud’ partitions. Regions of genome plasticity (RGPs) and spots of insertion were predicted (120) and subgraphs of the hotspots of interest generated by providing the sequence of the flanking proteins in a fasta file. The output of this PPanGGOLiN analysis and the code used to generate it are available at https://github.com/KLemonLab/DpiMGE_Manuscript/blob/master/SupplementalMethods_PPanGGOLiN.md. The subgraphs represented as inserts on Fig. 2B were obtained with the command ′ppanggolin align -p pangenome.h5 --getinfo --draw_related --proteins′ using the aa sequences for the proteins upstream and downstream of each spot of interest. Since PPanGGOLiN does not currently allow creation of subgraphs using GCs imported from external clustering methods, the pangenome was run again using the default PPanGGOLiN workflow with MMseqs2 clustering (default settings: --identity 0.8, --coverage 0.8 and --defrag).

### Characterization of mobile genetic elements (MGEs)

We searched all genomes for phage elements using the PHASTER database and web server (http://phaster.ca) on 11/8/2018 (56, 57). We took the “intact” phage elements as defined by a phage score of >90 and queried their ORFs using blastp to manually re-annotate their phage genes in the SnapGene GUI.

We searched for plasmid elements in all genomes using the PlasmidFinder 2.0 database and GUI interface (https://cge.cbs.dtu.dk/services/PlasmidFinder/) on 11/13/2018 following the default parameters (62). For strains with hits for a plasmid element, ORFs 1000 kb upstream and downstream of the element were queried through blastp. Manual gene re-annotation was done on the SnapGene GUI platform.

The summary file exported from the Anvi’o pangenomic analysis (see above) was also used for the identification of MGEs on the Prokka, COG20, Pfam and KOfam annotations. We identified 23 GCs as coding for putative transposases. GC alignments were visually inspected in AliView (121) and full-length representative sequences selected for Pfam search at the Pfam batch sequence search/HMMER website (112, 122). We identified 8 GCs with complete (≥80% coverage) Pfam Transposase (tnp) domains as true predicted transposases and 5 GCs with complete (≥80% coverage) Pfam rve domains as integrases. We used Bacterial Operon Finder for Functional Organization, aka BOFFO, which is described and available at https://github.com/FredHutch/boffo, to identify the gene neighborhoods in which the selected transposases and integrases were located across all 28 *D. pigrum* genomes (**File S2, Table A**). The approach used by BOFFO is to search for a set of defined query genes across a collection of reference genomes by translated amino acid alignment, and then to summarize the results by their physical colocation and organization. In this way, operon structure can be identified as the consistent colocation of a set of genes across multiple genomes in the same relative orientation (including both position and strand). The groups of genes identified with BOFFO at minimum percent identity 85% and minimum coverage 80% were visualized using clinker (https://github.com/gamcil/clinker) (123) and summary data provided in (**File S2, Table A**) was calculated using the matrixStats package (https://github.com/HenrikBengtsson/matrixStats). Detailed methods for this part of the analysis, as well as relevant files, are available at https://github.com/KLemonLab/DpiMGE_Manuscript/blob/master/SupplementalMethods_MGEs.md.

We similarly used BOFFO to identify the gene neighborhood of the group II intron identified with Anvi’o and PPanGGOLiN (GC_00000001). Using Pfam, we confirmed two predicted domains in a sequence from *D. pigrum* KPL3250 in GC_00000001—a reverse transcriptase and a maturase. The best hit in a blastx search with this same sequence against the Bacterial Group II Intron Database was to the bacterial class C intron-encoded protein from La.re.I1 in *Lactobacillus reuteri* with 44% identity and 65% similarity over 419 amino acids (70).

### Base modification analysis and prediction of restriction-modification systems

For methylome analysis, interpulse durations were measured and processed for all pulses aligned to each position in the reference sequence. We used Pacific Biosciences’ SMRTanalysis v8, which uses an *in silico* kinetic reference and a t-te st-based kinetic score detection of modified base positions, to identify modified positions (124).

We identified RM systems using SMRTseq data, as previously described (125), using the SEQWARE computer resource, a BLAST-based software module in combination with the curated restriction enzyme database (REBASE, http://rebase.neb.com/rebase/rebase.html) (71). Prediction was supported by sequence similarity, presence, and order of predictive functional motifs, plus the known genomic context and characteristics of empirically characterized RM system genes within REBASE. This facilitated reliable assignment of candidate methyltransferase genes to each modified motif based on their RM type.

### Detection of 5-methylcytosine

For *D. pigrum* CDC 4709-98 (aka KPL1934), the presence of 5-methylcytosine in the predicted methylation motif GCNGC was assessed as previously described (125). Briefly, gDNA harvested with the Masterpure Complete DNA/RNA Purification Kit was bisulfite treated using the EpiMark Bisulfite Conversion kit (NEB; Ipswich, MA, USA); both per manufacturer’s instructions, except for a final elution volume of 20 μL in the EpiMark kit. We then selected two genomic regions each ≤ 700 bp containing ≥ 4 GCNGC motifs. We PCR amplified each region from 1 μL of the converted gDNA using TaKaRa EpiTaq HS for bisulfite-treated DNA (Takara Bio USA; Mountain View, CA, USA) per manufacturer’s instructions with primers designed using MethPrimer: oKL732 (5’-AAGTTTATTTTTTTGAGTTTGTTG-3’), oKL733 (5’- TACCCATAAAATTATCACCTTC-3’), oKL734 (5’- ATTGATTTAGTAATTTTTTTGGAATAT-3’) and oKL735 (5’-TAAATAACTCTACAAAAAACTCAACTTACC-3’). After amplicon purification with the QIAquick PCR purification kit (final elution 40 μL; Qiagen; Germantown, MD, USA), we used Sanger sequencing (Macrogen, USA) of each PCR product to detect cytosine methylation within the predicted motif. Additional m5C-based modified motif analysis was carried out for *Dolosigranulum pigrum* KPL3250 using MFRE-Seq, as previously described (126).

### Prediction of CRISPR-Cas systems

CRISPR cas genes were detected using the CRISPRFinder (https://crispr.i2bc.paris-saclay.fr/Server/array) (127) and the array elements downstream from these genes were found using the CRISPRDetect software (crispr.otago.ac.nz/CRISPRDetect/predict_crispr_array.html) (73). The spacers identified using CRISPRDetect were queried through databases of possible phage targets in the Genbank-Phage, Refseq-Plasmid, and IMGVR databases with CRISPRtarget (bioanalysis.otago.ac.nz/CRISPRTarget/crispr_analysis.html) (73, 128), keeping hits with a cut-off score greater than 14. All gene and array element searches were completed on the webserver on 2/16/2019 using the default parameters. We also queried the genomes through CRISPRdb and CRISPRCompar (crispr.i2bc.paris-saclay.fr) website on 3/18/2019 to identify and annotate spacers shared among the different strains, keeping hits with scores higher than 15 to indicate similarity (127, 129, 130).

### Data Availability

All genomes are available in NCBI. **Table 1** lists the accession number for each *D. pigrum* strain genome used in this study.

## Acknowledgements

We are deeply grateful to the participants who donated nostril swabs samples at a 2017 and 2018 science festival. Their contribution was critical to expanding our knowledge of *D. pigrum.* We thank colleagues and lab members who provided invaluable assistance at both outreach events, in particular Javier Fernandez Juarez, Kerry Maguire, Pallavi Murugkar, Pooja Balani, Sowmya Balasubramanian, Fan Zhu, Andy Kaminsky, Andrew Collins, Brian Klein and Megan Lambert. For critical logistical support, we are grateful to Genevieve Holmes. We also thank Melinda M. Pettigrew and Yong Kong for advice on genome analysis as well as Tsute (George) Chen, Daniel Utter, Edoardo Pasolli, Nicola Segata, and Michael Wollenberg for their computational and phylogenetic advice over the course of the project. We thank members of the Johnston Lab, the KLemon Lab and the Starr-Johnston-Dewhirst-Lemon joint lab meeting for critique and suggestions.

## Authorship contributions

Conceptualization: KPL, SDB, CDJ. Methodology: SFR, SDB, IFE, CDJ, CAS, SLC, SME, THT, DSJ, SM, RJR. Strain isolation: SME, SFR, WG, SDB, LB, IFE. Investigation: SFR, CAS, SLC, SDB, SME, SM, CDJ, IFE. Interpretation of data: SFR, KPL, CDJ, IFE, SDB, CAS, SLC, SME, DSJ, SM, RJR. Visualization: SFR, CDJ, IFE. Wrote Original Draft: SFR, KPL, CDJ, IFE. Editing and review: SFR, IFE, CDJ, KPL, SDB, RJR, SME, THT, CAS, SM. All authors read and approved the final manuscript. Supervision: KPL, CDJ. Funding Acquisition: CDJ, KPL.

## Supplemental Materials

**File S1**. Supplemental Genomic Structural Analysis

**File S2**. Supplemental Genetic Elements and Defense Systems Analysis

**Table S1**. Pairwise SNP analysis

**Table S2**. RM Systems

**Table S3**. CRISPR-Cas systems

## Supplemental File S1

**Figure A.**
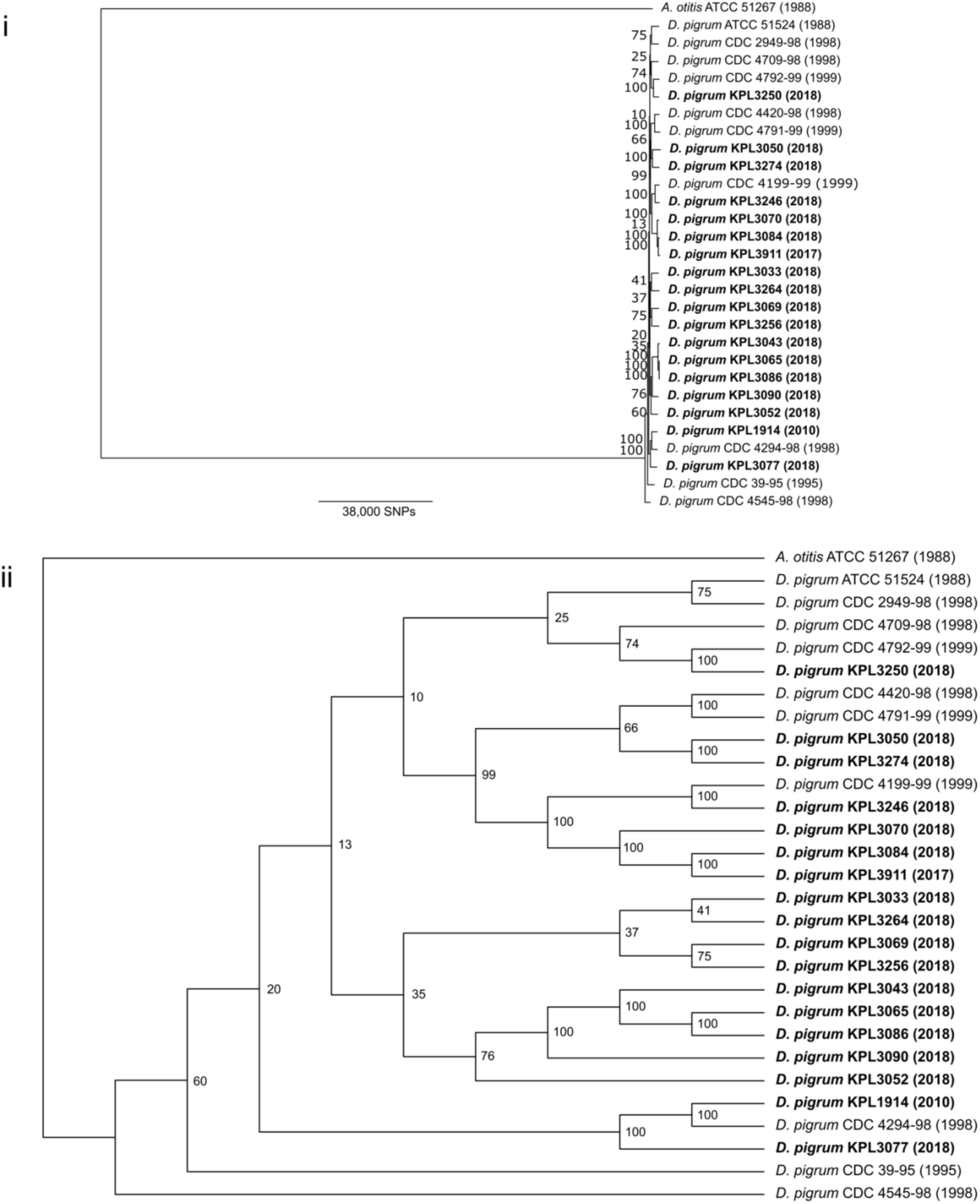
A core-gene phylogenomic tree including *Alloiococcus otitis* ATCC 51267 as an outgroup has poor resolution. (**i**) The tree with ancestral distance displayed. (**ii**) The tree as a cladogram. We generated a phylogenomic tree of 28 *D. pigrum* strains based on 789 concatenated core gene clusters (GCs) shared among *A. otitis* ATCC 51267 (NZ_JH992957) (1)—the most closely related genome-sequenced taxon to *D. pigrum* based on 16S rRNA gene sequence in both the Living Tree Project (2, 3) and the <u>eHOMD</u> (4)—and the 28 genomes with IQ-Tree using a GTR+F+R6 substitution model (BIC value 5743936.9087), 300 ML searches (ML=-2868528.1268) and 1000 ultrafast rapid Bootstraps. However, as shown above, the bootstrap values of the ancestral nodes for this tree were consistently low. There is also a deep branch separating *A. otitis* from all the *D. pigrum* strains. Repeated construction resulted in phylogenies with poorly supported nodes likely due to poor SNP resolution (Table S1), suggesting the need for a better outgroup for *D. pigrum*.

**Table A.** Pangenome contribution of all 28 *D. pigrum* genomes^a^.

| Strain name | Size (bp) | GC% | CDS | tRNA | rRNA | tmRNA | Core | Soft core | Shell | Cloud |
| --- | --- | --- | --- | --- | --- | --- | --- | --- | --- | --- |
| ATCC 51524 | 1,862,135 | 39.57% | 1684 | 23 | 9 | 1 | 1134 | 195 | 245 | 42 |
| KPL1914 | 1,890,316 | 39.53% | 1770 | 53 | 12 | 1 | 1134 | 197 | 293 | 52 |
| CDC 39-95 | 1,859,258 | 39.72% | 1708 | 52 | 6 | 1 | 1134 | 192 | 250 | 68 |
| CDC 2949-98 | 1,886,398 | 39.57% | 1693 | 53 | 6 | 1 | 1134 | 192 | 245 | 52 |
| CDC 4294-98 | 2,014,679 | 39.47% | 1914 | 55 | 5 | 1 | 1134 | 190 | 310 | 211 |
| CDC 4420-98 | 1,934,436 | 39.65% | 1776 | 55 | 4 | 1 | 1134 | 197 | 286 | 84 |
| CDC 4545-98 | 1,861,299 | 39.58% | 1704 | 53 | 4 | 1 | 1134 | 181 | 245 | 74 |
| CDC 4709-98 | 1,987,788 | 39.68% | 1778 | 54 | 12 | 2 | 1134 | 197 | 290 | 37 |
| CDC 4199-99 | 1,976,602 | 39.65% | 1795 | 56 | 7 | 1 | 1134 | 165 | 311 | 105 |
| CDC 4791-99 | 1,873,869 | 39.60% | 1692 | 54 | 5 | 1 | 1134 | 190 | 238 | 62 |
| CDC 4792-99 | 1,893,917 | 39.43% | 1734 | 56 | 6 | 1 | 1134 | 194 | 248 | 87 |
| KPL3033 | 1,958,196 | 39.55% | 1801 | 53 | 12 | 1 | 1134 | 196 | 321 | 72 |
| KPL3043 | 1,885,610 | 39.80% | 1714 | 53 | 12 | 1 | 1134 | 194 | 274 | 21 |
| KPL3050 | 2,043,806 | 39.30% | 1840 | 53 | 12 | 1 | 1134 | 193 | 339 | 72 |
| KPL3052 | 2,001,464 | 39.43% | 1820 | 53 | 12 | 1 | 1134 | 194 | 342 | 68 |
| KPL3065 | 1,878,426 | 39.82% | 1694 | 53 | 12 | 1 | 1134 | 197 | 278 | 13 |
| KPL3069 | 1,977,296 | 39.58% | 1870 | 53 | 12 | 1 | 1134 | 177 | 347 | 123 |
| KPL3070 | 1,883,862 | 39.76% | 1738 | 53 | 12 | 1 | 1134 | 192 | 292 | 36 |
| KPL3077 | 1,956,924 | 39.23% | 1838 | 53 | 12 | 1 | 1134 | 194 | 352 | 67 |
| KPL3084 | 1,906,822 | 39.61% | 1757 | 54 | 12 | 1 | 1134 | 193 | 302 | 36 |
| KPL3086 | 1,864,691 | 39.79% | 1706 | 53 | 12 | 1 | 1134 | 192 | 278 | 31 |
| KPL3090 | 2,009,643 | 39.74% | 1891 | 56 | 12 | 1 | 1134 | 193 | 310 | 127 |
| KPL3246 | 1,946,134 | 39.79% | 1720 | 54 | 12 | 1 | 1134 | 192 | 276 | 44 |
| KPL3250 | 1,904,333 | 39.61% | 1714 | 54 | 12 | 1 | 1134 | 196 | 263 | 44 |
| KPL3256 | 1,941,301 | 39.73% | 1800 | 54 | 12 | 1 | 1134 | 193 | 307 | 65 |
| KPL3264 | 1,984,655 | 39.70% | 1811 | 53 | 12 | 1 | 1134 | 186 | 323 | 86 |
| KPL3274 | 1,875,895 | 39.76% | 1684 | 53 | 12 | 1 | 1134 | 188 | 239 | 51 |
| KPL3911 | 1,900,145 | 39.59% | 1772 | 54 | 12 | 1 | 1134 | 193 | 304 | 44 |
| **AVERAGE** | **1927139** | **39.62%** | **1765** | **53** | **10** | **1** | **n.a.** | **191** | **290** | **67** |
| **MEDIAN** | **1905578** | **39.61%** | **1764** | **53** | **12** | **1** | **n.a.** | **193** | **291** | **64** |

<sup>a</sup>With addition of new strain genomes, the average genome size increased from 1.86 Mb (median 1.88 Mb)(5) to 1.93 Mb (median 1.91 Mb), while the GC content remained at 39.6%. The CDS, tRNA, rRNA, and tmRNA are based on the Prokka::prodigal annotations. The average length of the CDS was 950 bp, while the average number of
genes per kb was 0.916. There was also an increase in the number of predicted coding sequences from 1750 (median 1734) (5) to 1765 (median 1764). The predicted tRNA, rRNA, and tmRNA sequences averaged to 53, 10, and 1, respectively.

**Figure B.**
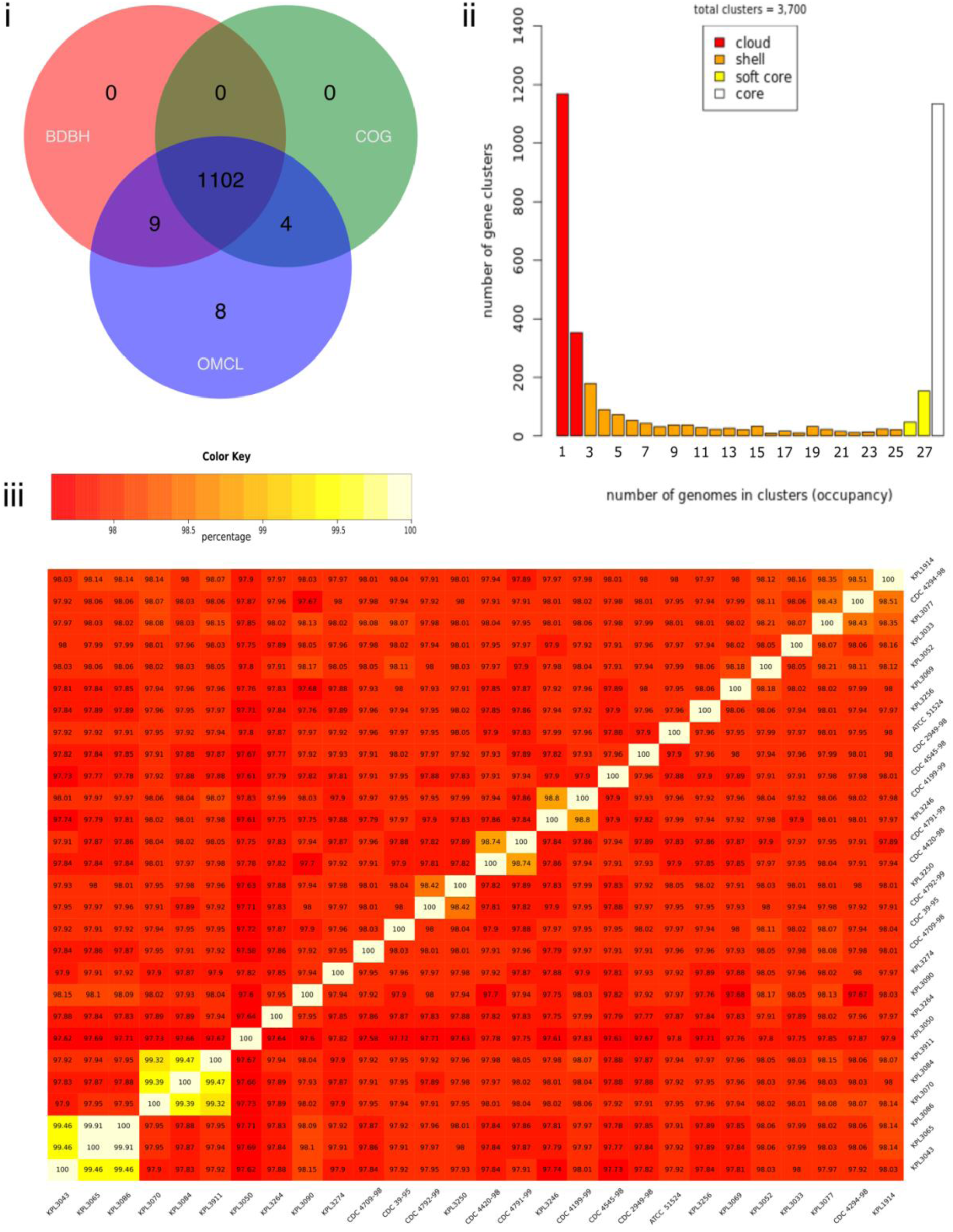
The conservative core genome of *D. pigrum* has 1102 GCs with a very high degree of nucleotide conservation among strains. (**i**) We used the intersection of the BDBH, COG triangle, and OMCL algorithm results to determine a conservative core of 1102 single-copy GCs using the 28 *D. pigrum* genomes on GET_HOMOLOGUES v. 3.1.4. (**ii**) We determined the GCs present in the core (n = 28), soft core (28 > n ≥ 26), shell (26 > n ≥ 3), and cloud (n ≤ 2) using the results from the OMCL and COG triangle algorithms. Of the 3700 genes clusters 30.6% are core genes (1134/3700), 5.41% are soft core (200/3700), 22.8% are shell (845/3700), and 41.1% are cloud (1521/3700). (**iii**) All paired strains in the pangenome share above 97.58% similarity at the nucleotide level. We based this on the average nucleotide identity of the homologues determined using the OMCL algorithm.

**Figure C.**
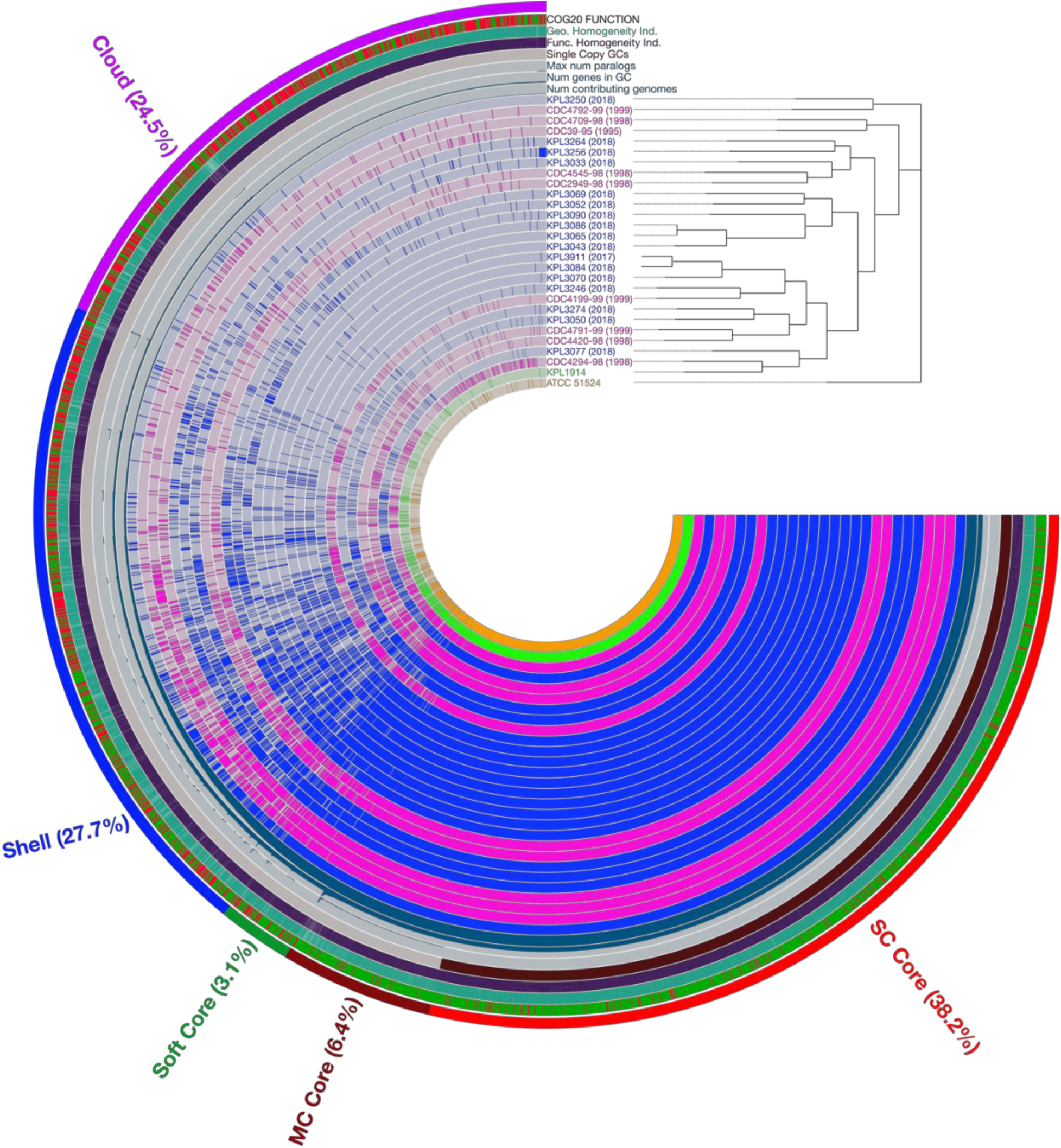
The *D. pigrum* Anvi’o pangenome is similar to the Get_Homologues pangenome. The biggest difference between the GET_HOMOLOGUES- and Anvi’o- defined pangenome was the migration of many of the GET_HOMOLOGUES cloud GCs to the Anvi’o multicopy core, causing the pangenome size to drop from 3,700 to 2,905 GCs. Occupancy of the gene clusters among the 28 genomes is displayed with strains collected in 2018 and 2017 (blue), strains from the CDC (pink), the ATCC strain (gold), and the strain collected in 2010 (green). Strains are arranged based on the phylogeny from Fig. 1 (right). With Anvi’o, 44.7% of the gene clusters (1,298/2,905) were in the core, 3.1% (90/2,905) in the soft core (green), 27.7% (805/2,905) in the shell (blue), and 24.5% (712/2,905) in the cloud (purple). A closer inspection of the core also showed 38.2% (1111/2,905) of the gene clusters are in the conservative single-copy (SC) core (red) and 6.4% (187/2,905) are in the multicopy (MC) core (dark red). The interactive Anvi’o pangenome can also be found on our GitHub (https://github.com/KLemonLab/DpiMGE_Manuscript/blob/master/SupplementalMethods_Anvio.md).

**Figure D.**
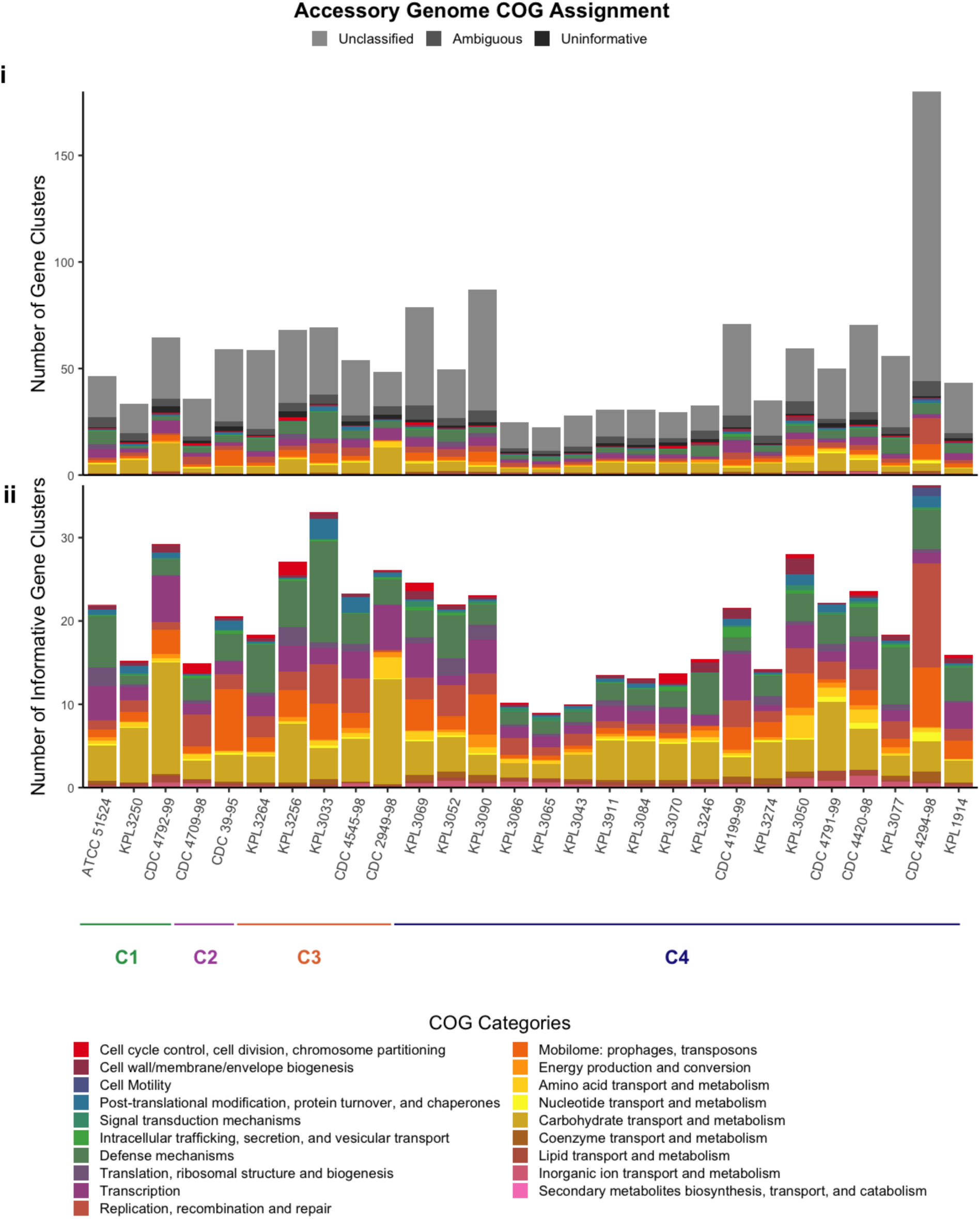
The accessory genome size varies but the functions are similar among the 28 *D. pigrum* genomes. **(i)** Using the functional annotations of the Accessory GCs identified by Anvi’o (defined as “shell” plus “cloud” bins), we found that most of the accessory GCs among all our genomes either had no annotation (“unclassified”, 53.6%), had an ambiguous COG categorization (“ambiguous”, 6.6%) or belong to the uninformative S or R COG category (“uninformative”, 2.6%). Only 37.2% of the gene clusters in the accessory genome had informative COG annotations (colored categories). **(ii)** Out of the informative gene clusters, there was a similar proportional distribution of accessory functions among all 28 genomes with a large fraction dedicated to ‘defense mechanisms’ (olive) and ‘carbohydrate transport and metabolism’ (khaki). The size of the accessory genomes varied among the strains with CDC 4294-98 and KPL3090 having the largest accessory size as compared to KPL3065 and KPL3086 with the smallest. Horizontal bars indicate clades: clade 1 green, clade 2 purple, clade 3 orange, and clade 4 blue.

## Supplemental File S2

### A 100-kb region with a higher frequency of indels

Closer inspection of the ∼650,000-750,000 position in the MAUVE alignment (**Fig. 2A; Fig. A**) showed the presence of three <u>s</u>ites containing sequencing <u>h</u>eterogeneity (HS for heterogeneity sites) bounded by >10 kb blocks of homology. The first site (HS1), between *abgT* (p-aminobenzoyl-glutamate transport protein) and *kynB* (kynurenine formamidase), contained a PTS (mannitol) system for metabolism. However, the extent of completeness of this PTS system varied among the *D. pigrum* isolates and was absent in KPL3274 and KPL3256. The next site (a few kb downstream of HS1), between *yxdM* (ABC transporter permease protein YxdM) and *ghrA* (glyoxylate/hydroxypyruvate reductase A), contained one of the two possible CRISPR-Cas systems, with all genomes having at least one type (CS1, **Fig. 8A-B**). Finally, the site (HS2), between *nth* (endonuclease III) and *clsA_1* (cardiolipin synthase A), variably harbored two types of aspartate and threonine synthase genes. One type in KPL3033 and KPL3052; none in KPL1914, KPL3077, and KPL3274; and a variation of the second type among the remaining genomes.

**Figure A.**
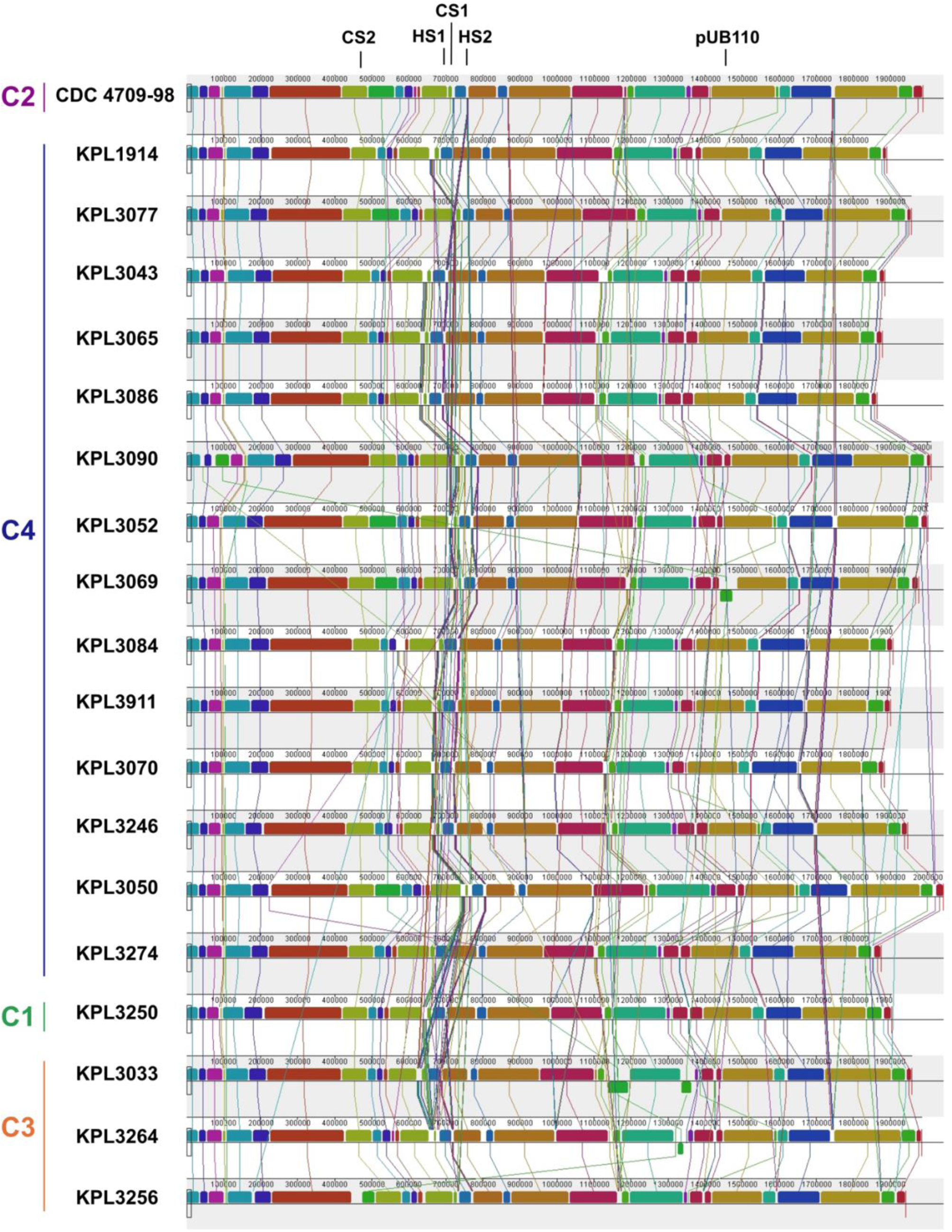
Other partial mobile genetic elements predicted and their locations. The MAUVE alignment with the locations of the partial pUB110 insertion, the two CRISPR-Cas insertion sites (CS1 and CS2) and two other sites of heterogeneity (HS1 and HS2) in a region with higher variability including CS1. Vertical bars represent clades: clade 1 green, clade 2 purple, clade 3 orange, and clade 4 blue.

### A region with a higher frequency of phage and plasmid integration events, including integrated plasmid sequence derived from pUB110

No autonomous plasmids were detected among the 28 genomes using the Gram-positive plasmid database, PlasmidFinder (1). However, we detected a sequence match to pUB110 (GenBank ID: NC_001384.1), a plasmid originally isolated from *S. aureus,* integrated in four of the *D. pigrum* isolates (KPL3043, KPL3065/KPL3086 in Clade 4 and CDC 4709-98 in Clade 2; **Fig. 1**). This included a perfect match (708/708 bp) to the *repB* gene of pUB110, which was located adjacent to three other genes that are also present in pUB110 and predicted to encode for a kanamycin nucleotidyltransferase ANT(4’)-lb (*knt*), a bleomycin resistance protein (*ble*), and a plasmid recombination protein (*pre-2*) (**Fig. B**). Inspection of the region adjacent to the genes from pUB110, at position ∼1.45Mb in the MAUVE alignment, showed evidence of plasmid elements present in 11 of the other closed *D. pigrum* genomes, suggesting this region has a higher frequency of phage and plasmid integration events. For example, a 73% (288/391 bp) match to a different (non-pUB110) *repB* gene present in plasmids of other closely related Firmicutes including *Lactobacillus plantarum* (KU707950) (2) and *Lactobacillus curvatus* (CP031007) (3). A subset of the genomes, including six from clade 4 (KPL3077, KPL3090, KPL3069, KPL3084, KPL3911, KPL3070) and two from clade 3 (KPL3264, and KPL3256), had an *ltrA* gene (group II reverse transcriptase/maturase) at this location suggesting it is a region with a high frequency of phage and plasmid integration events.

**Figure B.**
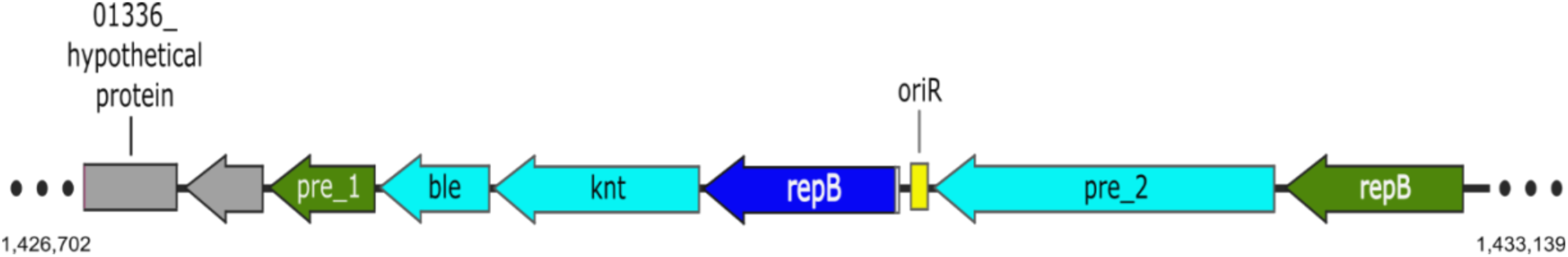
Genes corresponding to *S. aureus* plasmid pUB110 were present in 4 of 28 *D. pigrum* genomes. As illustrated with this map from *D. pigrum* KPL3043, four genes from plasmid pUB110 (*repB* in blue, remainder in cyan) were present in strains KPL3043, KPL3065, KPL3086, and CDC 4709-98: bleomycin resistance protein (*ble*), kanamycin nucleotidyltransferase ANT(4’)-lb (*knt*), replication protein B (*repB*), and plasmid recombination enzyme (*pre_2*).

### A subset of *D. pigrum* isolates encode for innate antibiotic resistance to kanamycin and/or erythromycin

Harmless members of human microbiota can serve as reservoirs of antibiotic resistance. Therefore, we searched for antibiotic resistance genes in these 28 *D. pigrum* genomes finding that six of the isolates encode predicted antibiotic resistance genes. Four genomes had a 100% identity match based on the RGI (4, 5) to a kanamycin nucleotidyltransferase ANT(4’)-lb that is encoded in integrated sequence homologous to pUB110 (CDC 4709-98, KPL3043, and KPL3065/KPL3086, the latter two having nearly identical genomes) (**Fig. B**). The latter three of these along with another clade 4 isolate (KPL3050) and a clade 1 isolate (KPL3250) also encoded an rRNA adenine N-6-methyltransferase (ErmT). With a strict identity match in the RGI of 86.53%, the predicted *ermT* gene is located within the CRISPR array of a subtype II-A system (**Fig. 8A-B**) in a different location in the genome than the kanamycin nucleotidyltransferase (CS1 vs. pUB110 in **Fig. A**).

### The four predicted *D. pigrum* prophage have few and dissimilar matches to known phages

We found four predicted mostly intact prophages in three of the *D. pigrum* genomes (**Fig. 5**) as defined by PHASTER’s scoring system (Intact: score > 90; Questionable: score 70-90; Incomplete: score < 70): L1 in KPL3069 (score of 150; C4); L4 in KPL3256 (score 130; C3); and L2 and L3 in KPL3090 (score 130 and 100, respectively; C4). CDS from prophages L1-4 displayed few and disparate matches to known phage genes. For example, prophage L1 had similarity to up to 30 different phage species among 56.3% (40/71) of its non-hypothetical proteins. The most common of these phage elements in L1 each only covered 11.3% (8/71) of the CDS in the phage region and were from a *Streptococcus pyogenes* host: temperate phage phiNIH1.1 (NC_003157) (6), and *Streptococcus* prophage 315.4 (NC_004587) (7). Temperate phage phiNIH1.1, which is integrated in the *Streptococcus pyogenes* M3/T3/subtype emm3.1 genome, is known for infecting lactic acid bacteria (6). Similarly, L4’s most common match to known phages only had a 9.3% (7/75) CDS identity match to the temperate *Enterococcus* phage EFC-1 from *Enterococcus faecalis* KBL101 (NC_025453) (8). The most common phage hits for L2 and L3 were to *Streptococcus* phage phi O1205 (NC_004303) (9) and *Streptococcus* phage SMP (NC_008721) (10) with a 7.6% (6/79) and 15.8% (12/76) CDS percent identity, respectively. *Streptococcus* phage SMP is integrated in the genome of a *Streptococcus suis* type 2 strain, which was isolated from the nasal swab of a healthy Bama pig (10).

**Table A.**
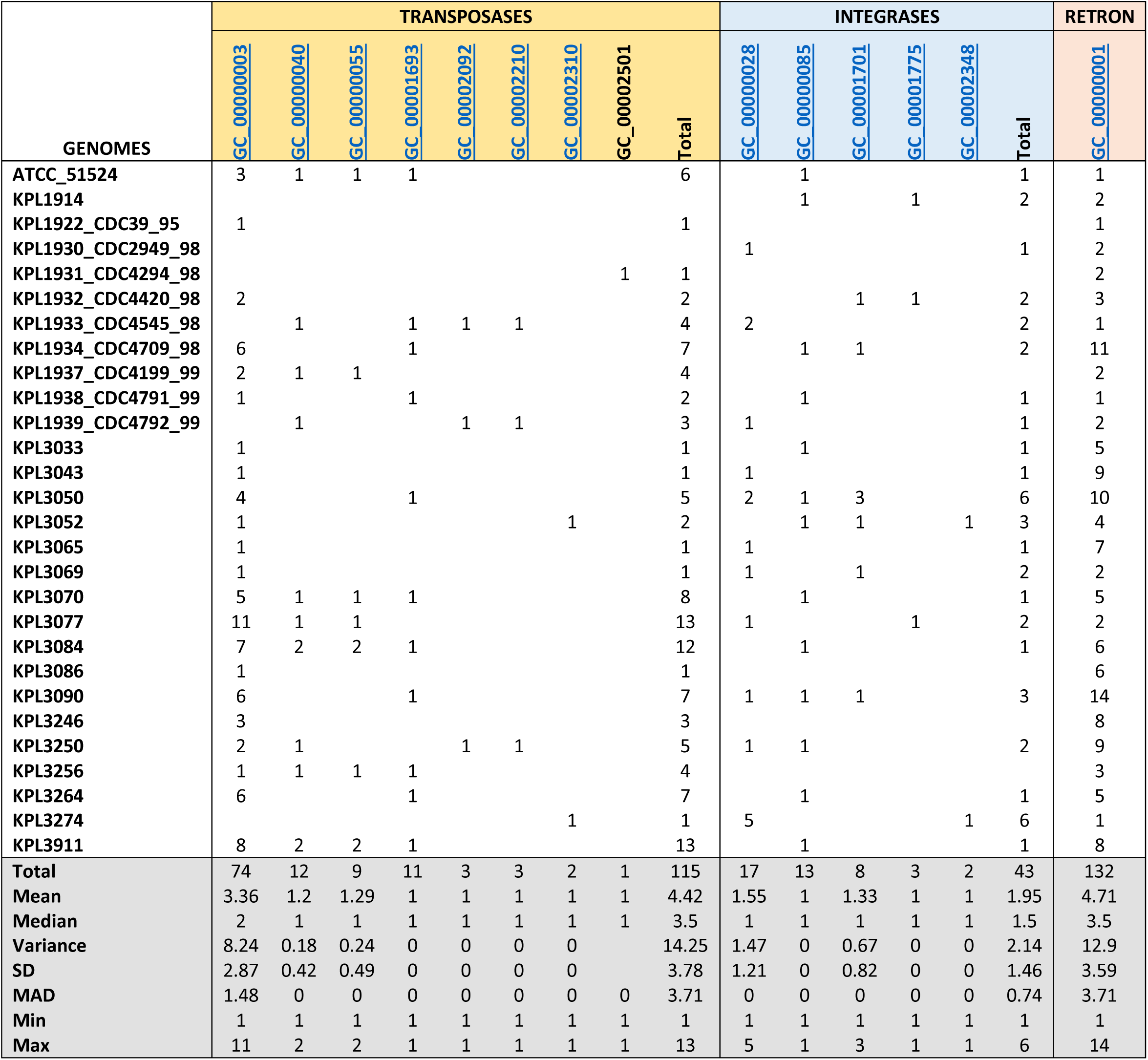
Predicted *D. pigrum* transposases, integrases and group II intron.

### Detailed description of *D. pigrum* restriction-modification (RM) systems

We identified Type I-IV RM systems across the 28 *D. pigrum* genomes located in three RMS insertion sites (**Fig. 7**). Type I RM systems typically consist of three separate genes encoding a restriction subunit (*hsdR*), a modification subunit (*hsdM*) and a recognition/specificity (*hsdS*) subunit. These three form a multi-subunit complex to catalyze both restriction and modification activities that generally target bipartite DNA motifs comprising two half-sequences separated by a gap (11). Two *D. pigrum* isolates, KPL3246 and KPL3264, were found to contain individual Type I systems and as each methylome contained characteristic bipartite motifs, CRTAN_7_TCNNC and CTAN_7_TGC respectively, associated with m6A modifications they were assigned as active.

Type II RM systems generally consist of independent restriction endonuclease (REase) and modification methyltransferase (MTase) proteins that do not form a complex, but instead recognize the same target motif and compete for activity. We identified 20 individual Type II systems across the 28 *D. pigrum* isolates and assigned the target motif to 15 of these based upon methylome analysis and/or REBASE homology to empirically characterized RM systems from other species (**Fig. 7A, Table S2**). A candidate motif was not detected for an m5C-associated RM system that co-occurred immediately downstream of the G^m5^CNGC system in four isolates (KPL3264, KPL3911, KPL3084, and KPL3070). Additionally, we were unable to unambiguously assign motifs to two Type IIG RM systems that occurred in KPL3256 (REBASE assignments: Dpi3256ORF5220 and Dpi3256ORF1810), although well-informed guesses could be made (CCAGT and GACAG, respectively). Type IIG systems are defined by the presence of a single polypeptide including an REase and an MTase domain that share a target recognition domain. A single candidate modified motif for one of these systems was identified during SMRTseq (CC^m6^AGT).

Type III RM systems consist of two genes (mod and res) encoding protein subunits that function either in DNA recognition and modification (MTase) or restriction (REase) activities. All Type III REases recognize asymmetrical DNA sequences and the modified DNA bears methyl groups on only one strand of the DNA recognition motif. We found three *D. pigrum* isolates (KPL3274, KPL3052 and KPL3070) harbored characteristic Type III systems and assigned these to specific hemi-methylated recognition motifs (CAACA, GTCAT, YACAG) detected during methylome analysis (**Table S2**). In such systems, the REase must interact with two copies of its nonpalindromic recognition sequence and the sites must be in an inverse orientation within the substrate DNA molecule for cleavage, which occurs at a specific distance away from one of the recognition sequences.

Type IV restriction enzymes are technically not true RM systems since they comprise only a restriction enzyme and have no accompanying methylase. These restriction enzymes recognize and cut only modified DNA, including methylated, hydroxymethylated and glucosyl-hydroxymethylated bases. We identified a largely conserved single gene Type IV system in 10 *D. pigrum* isolates (**Fig. 7; Table S2**). However, the targets of Type IV systems cannot be determined through SMRT sequencing and methylome analysis and, therefore, exact determination of its target recognition motif was not possible here. Nevertheless, there are three well characterized Type IV systems to date each with defined sequence preference and cleavage position (12). The *D. pigrum* Type IV system is most homologous (99% coverage/42% identity) to the SauUSI of *Staphylococcus aureus*; a modified cytosine restriction system targeting S^5m^CNGS (where S is C or G), but this level of homology is insufficient to confirm the exact modified motif targeted in the C. *pigrum* system (13).

**Figure C.**
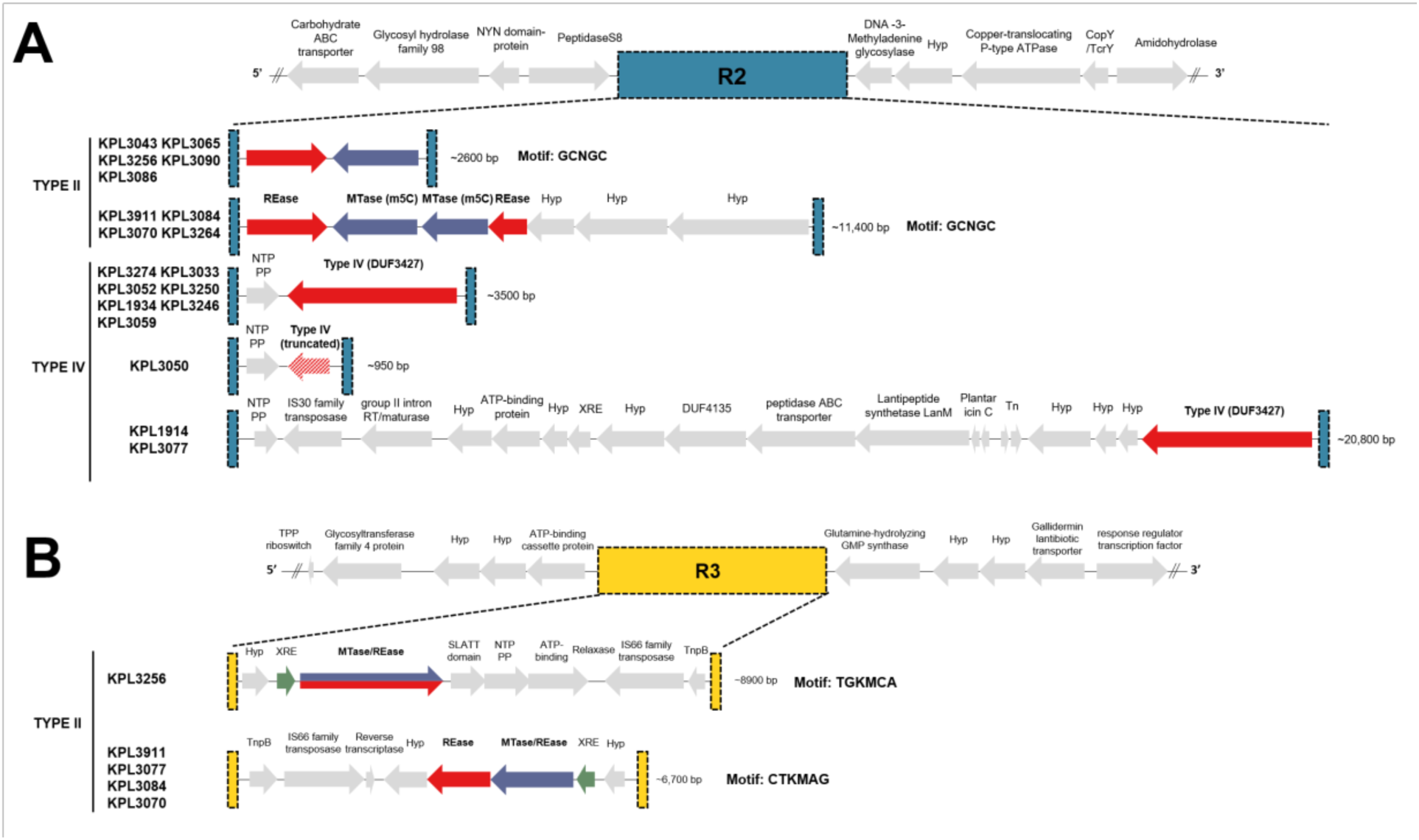
**A)** The organization of gene clusters within RMS integrative site 2 (R2), which primarily encodes a Type II m5C-associated RM system or a Type IV restriction system across the strain collection. R2 is flanked upstream by a region containing genes for a YacP-like NYN domain protein and an S8 family serine peptidase type protein, and downstream by genes for a DNA-3-methyladenine glycosylase protein, a hypothetical protein, and a copper-translocating P-type ATPase. **B)** The organization of gene clusters within integrative hotspot 3, which encodes two different Type II RM systems. Arrows represent the direction of translation and the relative sizes of open reading frames (ORFs). Putative control proteins are highlighted in green, hsdR and hsdM in red and blue, respectively, and fused hsdR/M ORFs are both red and blue. Proteins not identified as part of the R-M system or those with currently unknown function are shown in grey.

### *D. pigrum* CRISPR spacers have few matches to known MGEs

Two short-branched terminal clades (KPL3070, KPL3084, KPL3911 and KPL3043, KPL3065, KPL3086) shared the most spacers among all the isolates. However, only 9 of their spacers (Fig. 8B) had significant (cut-off score ≥ 15) matches to known MGEs. The KPL3070- containing clade had spacers with similarity to *Prochlorococcus* phage P-HM2 (GU075905; spacer 123) (14), *Lactobacillus plantarum* bacteriophage phiJL-1 (AY236756; spacer 131) (15), *Citrobacter* phage Moon (KM236240; spacer 136) (16), *Vibrio* phage 1.033.O._10N.222.49.B8 (MG592417; spacer 137) (17), *Pseudomonas sp.* Leaf58 plasmid pBASL58 (NZ_CP032678.1; spacer 142) (18), and the *Cupriavidus metallidurans* CH34 megaplasmid (NC_007974.2; spacer 143) (19). The KPL3043- containing clade had spacers with similarity to *Enterococcus* phage 156 (LR031359; spacer 7) (20), *Pectobacterium* phage CBB (KU574722; spacer 11) (21), and the *Bacillus* phage Page (KF669655; spacer 22) (22). Besides these two sets of closely related isolates, other less closely related isolates also shared more than three spacers. KPL3090 and KPL3052 both further apart in Clade 4 shared 7 spacers, which included matches to *Enterobacteria* phage RB49 (AY343333; spacer 39) (23) and the JP555 plasmid pJFP55H from *Clostridium perfringens* JP55 strain (NZ_CP013043.1; spacer 47) (24). All spacers sequences with their genome matches can be found in **Table S3B**.

## Supplemental methods

### Characterization of Virulence Factors

We found no matches when searching for predicted virulence factors among the 28 *D. pigrum* genomes by comparing to those encoded by *S. aureus* and *Enterococcus* using the VirulenceFinder2.0 online software (https://cge.cbs.dtu.dk/services/VirulenceFinder/) on 12/13/2020 (25).

### Characterization of innate antibiotic resistance genes

Prediction of antibiotic resistance was determined by querying the genomes in parallel through the Comprehensive Antibiotic Resistance Database (CARD) in the Resistance Gene Identifier (RGI, version 3.1.0) API platform using default settings (4, 5). Results were considered significant if either a perfect or strict match based on a protein homolog AMR detection model was detected.

## REFERENCES

1. Laufer AS, Metlay JP, Gent JF, Fennie KP, Kong Y, Pettigrew MM. 2011. Microbial communities of the upper respiratory tract and otitis media in children. mBio 2:e00245–10.

2. Pettigrew MM, Laufer AS, Gent JF, Kong Y, Fennie KP, Metlay JP. 2012. Upper respiratory tract microbial communities, acute otitis media pathogens, and antibiotic use in healthy and sick children. Appl Environ Microbiol 78:6262–70.

3. Biesbroek G, Bosch AA, Wang X, Keijser BJ, Veenhoven RH, Sanders EA, Bogaert D. 2014. The impact of breastfeeding on nasopharyngeal microbial communities in infants. American journal of respiratory and critical care medicine 190:298–308.

4. Biesbroek G, Tsivtsivadze E, Sanders EA, Montijn R, Veenhoven RH, Keijser BJ, Bogaert D. 2014. Early respiratory microbiota composition determines bacterial succession patterns and respiratory health in children. American journal of respiratory and critical care medicine 190:1283–92.

5. Liu CM, Price LB, Hungate BA, Abraham AG, Larsen LA, Christensen K, Stegger M, Skov R, Andersen PS. 2015. *Staphylococcus aureus* and the ecology of the nasal microbiome. Sci Adv 1:e1400216.

6. Teo SM, Mok D, Pham K, Kusel M, Serralha M, Troy N, Holt BJ, Hales BJ, Walker ML, Hollams E, Bochkov YA, Grindle K, Johnston SL, Gern JE, Sly PD, Holt PG, Holt KE, Inouye M. 2015. The infant nasopharyngeal microbiome impacts severity of lower respiratory infection and risk of asthma development. Cell Host Microbe 17:704–15.

7. Bosch A, Levin E, van Houten MA, Hasrat R, Kalkman G, Biesbroek G, de Steenhuijsen Piters WAA, de Groot PCM, Pernet P, Keijser BJF, Sanders EAM, Bogaert D. 2016. Development of Upper Respiratory Tract Microbiota in Infancy is Affected by Mode of Delivery. EBioMedicine 9:336–345.

8. Bomar L, Brugger SD, Yost BH, Davies SS, Lemon KP. 2016. *Corynebacterium accolens* Releases Antipneumococcal Free Fatty Acids from Human Nostril and Skin Surface Triacylglycerols. mBio 7:e01725–15.

9. Zhang M, Wang R, Liao Y, Buijs MJ, Li J. 2016. Profiling of Oral and Nasal Microbiome in Children With Cleft Palate. Cleft Palate Craniofac J 53:332–8.

10. Salter SJ, Turner C, Watthanaworawit W, de Goffau MC, Wagner J, Parkhill J, Bentley SD, Goldblatt D, Nosten F, Turner P. 2017. A longitudinal study of the infant nasopharyngeal microbiota: The effects of age, illness and antibiotic use in a cohort of South East Asian children. PLoS Negl Trop Dis 11:e0005975.

11. Bosch A, de Steenhuijsen Piters WAA, van Houten MA, Chu M, Biesbroek G, Kool J, Pernet P, de Groot PCM, Eijkemans MJC, Keijser BJF, Sanders EAM, Bogaert D. 2017. Maturation of the Infant Respiratory Microbiota, Environmental Drivers, and Health Consequences. A Prospective Cohort Study. Am J Respir Crit Care Med 196:1582–1590.

12. Kelly MS, Surette MG, Smieja M, Pernica JM, Rossi L, Luinstra K, Steenhoff AP, Feemster KA, Goldfarb DM, Arscott-Mills T, Boiditswe S, Rulaganyang I, Muthoga C, Gaofiwe L, Mazhani T, Rawls JF, Cunningham CK, Shah SS, Seed PC. 2017. The Nasopharyngeal Microbiota of Children With Respiratory Infections in Botswana. Pediatr Infect Dis J 36:e211–e218.

13. Hasegawa K, Linnemann RW, Mansbach JM, Ajami NJ, Espinola JA, Petrosino JF, Piedra PA, Stevenson MD, Sullivan AF, Thompson AD, Camargo CA, Jr. 2017. Nasal Airway Microbiota Profile and Severe Bronchiolitis in Infants: A Case-control Study. Pediatr Infect Dis J 36:1044–1051.

14. Langevin S, Pichon M, Smith E, Morrison J, Bent Z, Green R, Barker K, Solberg O, Gillet Y, Javouhey E, Lina B, Katze MG, Josset L. 2017. Early nasopharyngeal microbial signature associated with severe influenza in children: a retrospective pilot study. J Gen Virol 98:2425–2437.

15. Lappan R, Imbrogno K, Sikazwe C, Anderson D, Mok D, Coates H, Vijayasekaran S, Bumbak P, Blyth CC, Jamieson SE, Peacock CS. 2018. A microbiome case-control study of recurrent acute otitis media identified potentially protective bacterial genera. BMC Microbiol 18:13.

16. Escapa IF, Chen T, Huang Y, Gajare P, Dewhirst FE, Lemon KP. 2018. New Insights into Human Nostril Microbiome from the Expanded Human Oral Microbiome Database (eHOMD): a Resource for the Microbiome of the Human Aerodigestive Tract. mSystems 3.

17. Wen Z, Xie G, Zhou Q, Qiu C, Li J, Hu Q, Dai W, Li D, Zheng Y, Wen F. 2018. Distinct Nasopharyngeal and Oropharyngeal Microbiota of Children with Influenza A Virus Compared with Healthy Children. Biomed Res Int 2018:6362716.

18. Copeland E, Leonard K, Carney R, Kong J, Forer M, Naidoo Y, Oliver BGG, Seymour JR, Woodcock S, Burke CM, Stow NW. 2018. Chronic Rhinosinusitis: Potential Role of Microbial Dysbiosis and Recommendations for Sampling Sites. Front Cell Infect Microbiol 8:57.

19. Toivonen L, Hasegawa K, Waris M, Ajami NJ, Petrosino JF, Camargo CA, Jr., Peltola V. 2019. Early nasal microbiota and acute respiratory infections during the first years of life. Thorax 74:592–599.

20. Camelo-Castillo A, Henares D, Brotons P, Galiana A, Rodriguez JC, Mira A, Munoz-Almagro C. 2019. Nasopharyngeal Microbiota in Children With Invasive Pneumococcal Disease: Identification of Bacteria With Potential Disease-Promoting and Protective Effects. Front Microbiol 10:11.

21. Man WH, Clerc M, de Steenhuijsen Piters WAA, van Houten MA, Chu M, Kool J, Keijser BJF, Sanders EAM, Bogaert D. 2019. Loss of Microbial Topography between Oral and Nasopharyngeal Microbiota and Development of Respiratory Infections Early in Life. Am J Respir Crit Care Med doi:10.1164/rccm.201810-1993OC.

22. Man WH, van Houten MA, Merelle ME, Vlieger AM, Chu M, Jansen NJG, Sanders EAM, Bogaert D. 2019. Bacterial and viral respiratory tract microbiota and host characteristics in children with lower respiratory tract infections: a matched case-control study. Lancet Respir Med 7:417–426.

23. Man WH, van Dongen TMA, Venekamp RP, Pluimakers VG, Chu M, van Houten MA, Sanders EAM, Schilder AGM, Bogaert D. 2019. Respiratory Microbiota Predicts Clinical Disease Course of Acute Otorrhea in Children With Tympanostomy Tubes. Pediatr Infect Dis J 38:e116–e125.

24. Gan W, Yang F, Tang Y, Zhou D, Qing D, Hu J, Liu S, Liu F, Meng J. 2019. The difference in nasal bacterial microbiome diversity between chronic rhinosinusitis patients with polyps and a control population. Int Forum Allergy Rhinol doi:10.1002/alr.22297.

25. de Steenhuijsen Piters WAA, Jochems SP, Mitsi E, Rylance J, Pojar S, Nikolaou E, German EL, Holloway M, Carniel BF, Chu M, Arp K, Sanders EAM, Ferreira DM, Bogaert D. 2019. Interaction between the nasal microbiota and *S. pneumoniae* in the context of live-attenuated influenza vaccine. Nat Commun 10:2981.

26. De Boeck I, Wittouck S, Martens K, Claes J, Jorissen M, Steelant B, van den Broek MFL, Seys SF, Hellings PW, Vanderveken OM, Lebeer S. 2019. Anterior Nares Diversity and Pathobionts Represent Sinus Microbiome in Chronic Rhinosinusitis. mSphere 4.

27. Man WH, Scheltema NM, Clerc M, van Houten MA, Nibbelke EE, Achten NB, Arp K, Sanders EAM, Bont LJ, Bogaert D. 2020. Infant respiratory syncytial virus prophylaxis and nasopharyngeal microbiota until 6 years of life: a subanalysis of the MAKI randomised controlled trial. Lancet Respir Med doi:10.1016/S2213-2600(19)30470-9.

28. Brugger SD, Eslami SM, Pettigrew MM, Escapa IF, Henke MT, Kong Y, Lemon KP. 2020. *Dolosigranulum pigrum* Cooperation and Competition in Human Nasal Microbiota. mSphere 5.

29. Ortiz Moyano R, Raya Tonetti F, Tomokiyo M, Kanmani P, Vizoso-Pinto MG, Kim H, Quilodran-Vega S, Melnikov V, Alvarez S, Takahashi H, Kurata S, Kitazawa H, Villena J. 2020. The Ability of Respiratory Commensal Bacteria to Beneficially Modulate the Lung Innate Immune Response Is a Strain Dependent Characteristic. Microorganisms 8.

30. Coleman A, Bialasiewicz S, Marsh RL, Grahn Hakansson E, Cottrell K, Wood A, Jayasundara N, Ware RS, Zaugg J, Sidjabat HE, Adams J, Ferguson J, Brown M, Roos K, Cervin A. 2021. Upper Respiratory Microbiota in Relation to Ear and Nose Health Among Australian Aboriginal and Torres Strait Islander Children. J Pediatric Infect Dis Soc doi:10.1093/jpids/piaa141.

31. Brugger SD, Bomar L, Lemon KP. 2016. Commensal-Pathogen Interactions along the Human Nasal Passages. PLoS pathogens 12:e1005633.

32. Krismer B, Weidenmaier C, Zipperer A, Peschel A. 2017. The commensal lifestyle of *Staphylococcus aureus* and its interactions with the nasal microbiota. Nat Rev Microbiol 15:675–687.

33. Man WH, de Steenhuijsen Piters WA, Bogaert D. 2017. The microbiota of the respiratory tract: gatekeeper to respiratory health. Nat Rev Microbiol 15:259–270.

34. Bomar L, Brugger SD, Lemon KP. 2018. Bacterial microbiota of the nasal passages across the span of human life. Curr Opin Microbiol 41:8–14.

35. Esposito S, Principi N. 2018. Impact of nasopharyngeal microbiota on the development of respiratory tract diseases. Eur J Clin Microbiol Infect Dis 37:1–7.

36. Mittal R, Sanchez-Luege SV, Wagner SM, Yan D, Liu XZ. 2019. Recent Perspectives on Gene-Microbe Interactions Determining Predisposition to Otitis Media. Front Genet 10:1230.

37. Yan M, Pamp SJ, Fukuyama J, Hwang PH, Cho DY, Holmes S, Relman DA. 2013. Nasal microenvironments and interspecific interactions influence nasal microbiota complexity and *S. aureus* carriage. Cell host & microbe 14:631–40.

38. Accorsi EK, Franzosa EA, Hsu T, Joice Cordy R, Maayan-Metzger A, Jaber H, Reiss-Mandel A, Kline M, DuLong C, Lipsitch M, Regev-Yochay G, Huttenhower C. 2020. Determinants of *Staphylococcus aureus* carriage in the developing infant nasal microbiome. Genome Biol 21:301.

39. Khamash DF, Mongodin EF, White JR, Voskertchian A, Hittle L, Colantuoni E, Milstone AM. 2019. The Association Between the Developing Nasal Microbiota of Hospitalized Neonates and Staphylococcus aureus Colonization. Open Forum Infect Dis 6:ofz062.

40. De Boeck I, Spacova I, Vanderveken OM, Lebeer S. 2021. Lactic acid bacteria as probiotics for the nose? Microb Biotechnol doi:10.1111/1751-7915.13759.

41. Tettelin H, Masignani V, Cieslewicz MJ, Donati C, Medini D, Ward NL, Angiuoli SV, Crabtree J, Jones AL, Durkin AS, Deboy RT, Davidsen TM, Mora M, Scarselli M, Margarit y Ros I, Peterson JD, Hauser CR, Sundaram JP, Nelson WC, Madupu R, Brinkac LM, Dodson RJ, Rosovitz MJ, Sullivan SA, Daugherty SC, Haft DH, Selengut J, Gwinn ML, Zhou L, Zafar N, Khouri H, Radune D, Dimitrov G, Watkins K, O’Connor KJ, Smith S, Utterback TR, White O, Rubens CE, Grandi G, Madoff LC, Kasper DL, Telford JL, Wessels MR, Rappuoli R, Fraser CM. 2005. Genome analysis of multiple pathogenic isolates of Streptococcus agalactiae: implications for the microbial “pan-genome”. Proc Natl Acad Sci U S A 102:13950–5.

42. Contreras-Moreira B, Vinuesa P. 2013. GET_HOMOLOGUES, a versatile software package for scalable and robust microbial pangenome analysis. Appl Environ Microbiol 79:7696–701.

43. Doron S, Melamed S, Ofir G, Leavitt A, Lopatina A, Keren M, Amitai G, Sorek R. 2018. Systematic discovery of antiphage defense systems in the microbial pangenome. Science 359.

44. Goldfarb T, Sberro H, Weinstock E, Cohen O, Doron S, Charpak-Amikam Y, Afik S, Ofir G, Sorek R. 2015. BREX is a novel phage resistance system widespread in microbial genomes. EMBO J 34:169–83.

45. Wang L, Jiang S, Deng Z, Dedon PC, Chen S. 2019. DNA phosphorothioate modification-a new multi-functional epigenetic system in bacteria. FEMS Microbiol Rev 43:109–122.

46. Horvath P, Barrangou R. 2010. CRISPR/Cas, the immune system of bacteria and archaea. Science 327:167–70.

47. Makarova KS, Wolf YI, Snir S, Koonin EV. 2011. Defense islands in bacterial and archaeal genomes and prediction of novel defense systems. J Bacteriol 193:6039–56.

48. Laclaire L, Facklam R. 2000. Antimicrobial susceptibility and clinical sources of *Dolosigranulum pigrum* cultures. Antimicrob Agents Chemother 44:2001–3.

49. Aguirre M, Collins MD. 1992. Phylogenetic analysis of *Alloiococcus otitis* gen. nov., sp. nov., an organism from human middle ear fluid. Int J Syst Bacteriol 42:79–83.

50. Darling AC, Mau B, Blattner FR, Perna NT. 2004. Mauve: multiple alignment of conserved genomic sequence with rearrangements. Genome Res 14:1394–403.

51. Darling AE, Mau B, Perna NT. 2010. progressiveMauve: multiple genome alignment with gene gain, loss and rearrangement. PLoS One 5:e11147.

52. Eren AM, Esen OC, Quince C, Vineis JH, Morrison HG, Sogin ML, Delmont TO. 2015. Anvi’o: an advanced analysis and visualization platform for ’omics data. PeerJ 3:e1319.

53. Delmont TO, Eren AM. 2018. Linking pangenomes and metagenomes: the Prochlorococcus metapangenome. PeerJ 6:e4320.

54. Tatusov RL, Fedorova ND, Jackson JD, Jacobs AR, Kiryutin B, Koonin EV, Krylov DM, Mazumder R, Mekhedov SL, Nikolskaya AN, Rao BS, Smirnov S, Sverdlov AV, Vasudevan S, Wolf YI, Yin JJ, Natale DA. 2003. The COG database: an updated version includes eukaryotes. BMC Bioinformatics 4:41.

55. Galperin MY, Makarova KS, Wolf YI, Koonin EV. 2015. Expanded microbial genome coverage and improved protein family annotation in the COG database. Nucleic Acids Research 43:D261–D269.

56. Zhou Y, Liang Y, Lynch KH, Dennis JJ, Wishart DS. 2011. PHAST: A Fast Phage Search Tool. Nucleic Acids Research 39:W347–W352.

57. Arndt D, Grant JR, Marcu A, Sajed T, Pon A, Liang Y, Wishart DS. 2016. PHASTER: a better, faster version of the PHAST phage search tool. Nucleic Acids Research 44:W16–W21.

58. Dorscht J, Klumpp J, Bielmann R, Schmelcher M, Born Y, Zimmer M, Calendar R, Loessner MJ. 2009. Comparative Genome Analysis of Listeria Bacteriophages Reveals Extensive Mosaicism, Programmed Translational Frameshifting, and a Novel Prophage Insertion Site. Journal of Bacteriology 191:7206–7215.

59. Zimmer M, Sattelberger E, Inman RB, Calendar R, Loessner MJ. 2003. Genome and proteome of Listeria monocytogenes phage PSA: an unusual case for programmed + 1 translational frameshifting in structural protein synthesis. Molecular Microbiology 50:303–317.

60. van Sinderen D, Karsens H, Kok J, Terpstra P, Ruiters MHJ, Venema G, Nauta A. 1996. Sequence analysis and molecular characterization of the temperate lactococcal bacteriophage r1t. Molecular Microbiology 19:1343–1355.

61. Beres SB, Sylva GL, Barbian KD, Lei B, Hoff JS, Mammarella ND, Liu M-Y, Smoot JC, Porcella SF, Parkins LD, Campbell DS, Smith TM, McCormick JK, Leung DYM, Schlievert PM, Musser JM. 2002. Genome sequence of a serotype M3 strain of group A *Streptococcus*: Phage-encoded toxins, the high-virulence phenotype, and clone emergence. Proceedings of the National Academy of Sciences 99:10078.

62. Carattoli A, Zankari E, García-Fernández A, Voldby Larsen M, Lund O, Villa L, Møller Aarestrup F, Hasman H. 2014. In SilicoDetection and Typing of Plasmids using PlasmidFinder and Plasmid Multilocus Sequence Typing. Antimicrobial Agents and Chemotherapy 58:3895–3903.

63. McArthur AG, Waglechner N, Nizam F, Yan A, Azad MA, Baylay AJ, Bhullar K, Canova MJ, De Pascale G, Ejim L, Kalan L, King AM, Koteva K, Morar M, Mulvey MR, O’Brien JS, Pawlowski AC, Piddock LJV, Spanogiannopoulos P, Sutherland AD, Tang I, Taylor PL, Thaker M, Wang W, Yan M, Yu T, Wright GD. 2013. The Comprehensive Antibiotic Resistance Database. Antimicrobial Agents and Chemotherapy 57:3348–3357.

64. Jia B, Raphenya AR, Alcock B, Waglechner N, Guo P, Tsang KK, Lago BA, Dave BM, Pereira S, Sharma AN, Doshi S, Courtot M, Lo R, Williams LE, Frye JG, Elsayegh T, Sardar D, Westman EL, Pawlowski AC, Johnson TA, Brinkman FSL, Wright GD, McArthur AG. 2017. CARD 2017: expansion and model-centric curation of the comprehensive antibiotic resistance database. Nucleic Acids Research 45:D566–D573.

65. Siguier P, Gourbeyre E, Chandler M. 2014. Bacterial insertion sequences: their genomic impact and diversity. FEMS Microbiol Rev 38:865–91.

66. Siguier P, Gourbeyre E, Varani A, Ton-Hoang B, Chandler M. 2015. Everyman’s Guide to Bacterial Insertion Sequences. Microbiol Spectr 3:MDNA3-0030-2014.

67. Aziz RK, Breitbart M, Edwards RA. 2010. Transposases are the most abundant, most ubiquitous genes in nature. Nucleic Acids Res 38:4207–17.

68. Gautreau G, Bazin A, Gachet M, Planel R, Burlot L, Dubois M, Perrin A, Medigue C, Calteau A, Cruveiller S, Matias C, Ambroise C, Rocha EPC, Vallenet D. 2020. PPanGGOLiN: Depicting microbial diversity via a partitioned pangenome graph. PLoS Comput Biol 16:e1007732.

69. McNeil BA, Semper C, Zimmerly S. 2016. Group II introns: versatile ribozymes and retroelements. Wiley Interdiscip Rev RNA 7:341–55.

70. Candales MA, Duong A, Hood KS, Li T, Neufeld RA, Sun R, McNeil BA, Wu L, Jarding AM, Zimmerly S. 2012. Database for bacterial group II introns. Nucleic Acids Res 40:D187–90.

71. Roberts RJ, Vincze T, Posfai J, Macelis D. 2015. REBASE--a database for DNA restriction and modification: enzymes, genes and genomes. Nucleic Acids Res 43:D298–9.

72. Koonin EV, Makarova KS, Wolf YI, Krupovic M. 2020. Evolutionary entanglement of mobile genetic elements and host defence systems: guns for hire. Nat Rev Genet 21:119–131.

73. Biswas A, Staals RHJ, Morales SE, Fineran PC, Brown CM. 2016. CRISPRDetect: A flexible algorithm to define CRISPR arrays. 17.

74. Bernheim A, Bikard D, Touchon M, Rocha EPC. 2020. Atypical organizations and epistatic interactions of CRISPRs and cas clusters in genomes and their mobile genetic elements. Nucleic Acids Res 48:748–760.

75. Crawley AB, Henriksen ED, Stout E, Brandt K, Barrangou R. 2018. Characterizing the activity of abundant, diverse and active CRISPR-Cas systems in lactobacilli. Scientific Reports 8:11544.

76. Hargreaves KR, Flores CO, Lawley TD, Clokie MRJ. 2014. Abundant and Diverse Clustered Regularly Interspaced Short Palindromic Repeat Spacers in Clostridium difficile Strains and Prophages Target Multiple Phage Types within This Pathogen. mBio 5:e01045–13-e0104.

77. Hall GS, Gordon S, Schroeder S, Smith K, Anthony K, Procop GW. 2001. Case of synovitis potentially caused by *Dolosigranulum pigrum*. J Clin Microbiol 39:1202–3.

78. Hoedemaekers A, Schulin T, Tonk B, Melchers WJ, Sturm PD. 2006. Ventilator-associated pneumonia caused by *Dolosigranulum pigrum*. J Clin Microbiol 44:3461–2.

79. Lin JC, Hou SJ, Huang LU, Sun JR, Chang WK, Lu JJ. 2006. Acute cholecystitis accompanied by acute pancreatitis potentially caused by *Dolosigranulum pigrum*. J Clin Microbiol 44:2298–9.

80. Lecuyer H, Audibert J, Bobigny A, Eckert C, Janniere-Nartey C, Buu-Hoi A, Mainardi JL, Podglajen I. 2007. *Dolosigranulum pigrum* causing nosocomial pneumonia and septicemia. J Clin Microbiol 45:3474–5.

81. Johnsen BO, Ronning EJ, Onken A, Figved W, Jenum PA. 2011. *Dolosigranulum pigrum* causing biomaterial-associated arthritis. APMIS 119:85–7.

82. Sherret J, Gajjar B, Ibrahim L, Mohamed Ahmed A, Panta UR. 2020. Dolosigranulum pigrum: Predicting Severity of Infection. Cureus 12:e9770.

83. Sampo M, Ghazouani O, Cadiou D, Trichet E, Hoffart L, Drancourt M. 2013. *Dolosigranulum pigrum* keratitis: a three-case series. BMC Ophthalmol 13:31.

84. Haas W, Gearinger LS, Hesje CK, Sanfilippo CM, Morris TW. 2012. Microbiological etiology and susceptibility of bacterial conjunctivitis isolates from clinical trials with ophthalmic, twice-daily besifloxacin. Adv Ther 29:442–55.

85. Venkateswaran N, Kalsow CM, Hindman HB. 2014. Phlyctenular keratoconjunctivitis associated with *Dolosigranulum pigrum*. Ocul Immunol Inflamm 22:242–5.

86. Monera-Lucas CE, Tarazona-Jaimes CP, Escolano-Serrano J, Martinez-Toldos JJ. 2020. Bilateral keratitis secondary to Dolosigranulum pigrum infection in a patient with HIV Infection. Enferm Infecc Microbiol Clin doi:10.1016/j.eimc.2020.10.017.

87. Oliveira PH, Touchon M, Cury J, Rocha EPC. 2017. The chromosomal organization of horizontal gene transfer in bacteria. Nat Commun 8:841.

88. McInerney JO, McNally A, O’Connell MJ. 2017. Why prokaryotes have pangenomes. Nat Microbiol 2:17040.

89. Lacks SA, Mannarelli BM, Springhorn SS, Greenberg B. 1986. Genetic basis of the complementary DpnI and DpnII restriction systems of S. pneumoniae: an intercellular cassette mechanism. Cell 46:993–1000.

90. Bondy-Denomy J, Davidson AR. 2014. To acquire or resist: the complex biological effects of CRISPR-Cas systems. Trends Microbiol 22:218–25.

91. Sanozky-Dawes R, Selle K, Klaenhammer T, O’Flaherty S, Barrangou R. 2015. Occurrence and activity of a type II CRISPR-Cas system in Lactobacillus gasseri. 161:1752–1761.

92. Shmakov SA, Sitnik V, Makarova KS, Wolf YI, Severinov KV, Koonin EV. 2017. The CRISPR Spacer Space Is Dominated by Sequences from Species-Specific Mobilomes. mBio 8.

93. EFSA Panel EFSA. 2012. Guidance on the assessment of bacterial susceptibility to antimicrobials of human and veterinary importance. EFSA Journal 10:2740.

94. Hunt M, Silva ND, Otto TD, Parkhill J, Keane JA, Harris SR. 2015. Circlator: automated circularization of genome assemblies using long sequencing reads. Genome Biology 16.

95. Tatusova T, DiCuccio M, Badretdin A, Chetvernin V, Nawrocki EP, Zaslavsky L, Lomsadze A, Pruitt KD, Borodovsky M, Ostell J. 2016. NCBI prokaryotic genome annotation pipeline. Nucleic Acids Res 44:6614–24.

96. Haft DH, DiCuccio M, Badretdin A, Brover V, Chetvernin V, O’Neill K, Li W, Chitsaz F, Derbyshire MK, Gonzales NR, Gwadz M, Lu F, Marchler GH, Song JS, Thanki N, Yamashita RA, Zheng C, Thibaud-Nissen F, Geer LY, Marchler-Bauer A, Pruitt KD. 2018. RefSeq: an update on prokaryotic genome annotation and curation. Nucleic Acids Res 46:D851–D860.

97. Seemann T. 2014. Prokka: rapid prokaryotic genome annotation. Bioinformatics 30:2068–9.

98. Vinuesa P, Contreras-Moreira B. 2015. Robust Identification of Orthologues and Paralogues for Microbial Pan-Genomics Using GET_HOMOLOGUES: A Case Study of pIncA/C Plasmids, p 203–232 doi:10.1007/978-1-4939-1720-4_14. Springer New York.

99. Kristensen DM, Kannan L, Coleman MK, Wolf YI, Sorokin A, Koonin EV, Mushegian A. 2010. A low-polynomial algorithm for assembling clusters of orthologous groups from intergenomic symmetric best matches. Bioinformatics 26:1481–1487.

100. Li L. 2003. OrthoMCL: Identification of Ortholog Groups for Eukaryotic Genomes. Genome Research 13:2178–2189.

101. Katoh K, Standley DM. 2013. MAFFT Multiple Sequence Alignment Software Version 7: Improvements in Performance and Usability. Molecular Biology and Evolution 30:772–780.

102. Galtier N, Gouy M, Gautier C. 1996. SEAVIEW and PHYLO_WIN: two graphic tools for sequence alignment and molecular phylogeny. 12:543–548.

103. Nguyen L-T, Schmidt HA, Von Haeseler A, Minh BQ. 2015. IQ-TREE: A Fast and Effective Stochastic Algorithm for Estimating Maximum-Likelihood Phylogenies. Molecular Biology and Evolution 32:268–274.

104. Kalyaanamoorthy S, Minh BQ, Wong TKF, Von Haeseler A, Jermiin LS. 2017. ModelFinder: fast model selection for accurate phylogenetic estimates. Nature Methods 14:587–589.

105. Hoang DT, Chernomor O, von Haeseler A, Minh BQ, Vinh LS. 2018. UFBoot2: Improving the Ultrafast Bootstrap Approximation. Mol Biol Evol 35:518–522.

106. Hyatt D, Chen G-L, Locascio PF, Land ML, Larimer FW, Hauser LJ. 2010. Prodigal: prokaryotic gene recognition and translation initiation site identification. BMC Bioinformatics 11:119.

107. Buchfink B, Xie C, Huson DH. 2015. Fast and sensitive protein alignment using DIAMOND. Nat Methods 12:59–60.

108. Tatusov RL, Koonin EV, Lipman DJ. 1997. A genomic perspective on protein families. Science 278:631–7.

109. Galperin MY, Wolf YI, Makarova KS, Vera Alvarez R, Landsman D, Koonin EV. 2021. COG database update: focus on microbial diversity, model organisms, and widespread pathogens. Nucleic Acids Res 49:D274–D281.

110. Kanehisa M, Goto S. 2000. KEGG: kyoto encyclopedia of genes and genomes. Nucleic Acids Res 28:27–30.

111. Kanehisa M, Sato Y, Kawashima M, Furumichi M, Tanabe M. 2016. KEGG as a reference resource for gene and protein annotation. Nucleic Acids Res 44:D457–62.

112. Mistry J, Chuguransky S, Williams L, Qureshi M, Salazar GA, Sonnhammer ELL, Tosatto SCE, Paladin L, Raj S, Richardson LJ, Finn RD, Bateman A. 2021. Pfam: The protein families database in 2021. Nucleic Acids Res 49:D412–D419.

113. Eddy SR. 2011. Accelerated Profile HMM Searches. PLoS Comput Biol 7:e1002195.

114. Altschul SF, Gish W, Miller W, Myers EW, Lipman DJ. 1990. Basic local alignment search tool. J Mol Biol 215:403–10.

115. Edgar RC. 2004. MUSCLE: multiple sequence alignment with high accuracy and high throughput. Nucleic Acids Res 32:1792–7.

116. Benedict MN, Henriksen JR, Metcalf WW, Whitaker RJ, Price ND. 2014. ITEP: an integrated toolkit for exploration of microbial pan-genomes. BMC Genomics 15:8.

117. van Dongen S, Abreu-Goodger C. 2012. Using MCL to Extract Clusters from Networks. *In* van Helden J, Toussaint A, Thieffry D (ed), Bacterial Molecular Networks Methods in Molecular Biology (Methods and Protocols), vol 804. Springer, New York, NY.

118. . R-Core-Team. 2020. R: A language and environment for statistical computing., R Foundation for Statistical Computing, Vienna, Austria. https://www.R-project.org/.

119. RStudio-Team. 2020. RStudio: Integrated Development for R. RStudio, PBC, Boston, MA. http://www.rstudio.com/.

120. Bazin A, Gautreau G, Médigue C, Vallenet D, Calteau A. 2020. panRGP: a pangenome-based method to predict genomic islands and explore their diversity. bioRxiv doi:10.1101/2020.03.26.007484:2020.03.26.007484.

121. Larsson A. 2014. AliView: a fast and lightweight alignment viewer and editor for large datasets. Bioinformatics 30:3276–8.

122. Potter SC, Luciani A, Eddy SR, Park Y, Lopez R, Finn RD. 2018. HMMER web server: 2018 update. Nucleic Acids Res 46:W200–W204.

123. Gilchrist CLM, Chooi YH. 2021. Clinker & clustermap.js: Automatic generation of gene cluster comparison figures. Bioinformatics doi:10.1093/bioinformatics/btab007.

124. Flusberg BA, Webster DR, Lee JH, Travers KJ, Olivares EC, Clark TA, Korlach J, Turner SW. 2010. Direct detection of DNA methylation during single-molecule, real-time sequencing. Nat Methods 7:461–5.

125. Johnston CD, Skeete CA, Fomenkov A, Roberts RJ, Rittling SR. 2017. Restriction-modification mediated barriers to exogenous DNA uptake and incorporation employed by Prevotella intermedia. PLoS One 12:e0185234.

126. Anton BP, Fomenkov A, Wu V, Roberts RJ. 2021. Genome-Wide Identification of 5-Methylcytosine Sites in Bacterial Genomes By High-Throughput Sequencing of MspJI Restriction Fragments. bioRxiv doi:10.1101/2021.02.10.430591:2021.02.10.430591.

127. Grissa I, Vergnaud G, Pourcel C. 2007. CRISPRFinder: a web tool to identify clustered regularly interspaced short palindromic repeats. 35:W52–W57.

128. Biswas A, Gagnon JN, Brouns SJJ, Fineran PC, Brown CM. 2013. CRISPRTarget. 10:817–827.

129. Grissa I, Vergnaud G, Pourcel C. 2007. The CRISPRdb database and tools to display CRISPRs and to generate dictionaries of spacers and repeats. BMC Bioinformatics 8:172.

130. Grissa I, Vergnaud G, Pourcel C. 2008. CRISPRcompar: a website to compare clustered regularly interspaced short palindromic repeats. 36:W145–W148.

131. Bastian M, Heymann S, Jacomy M. Gephi: An Open Source Software for Exploring and Manipulating Networks.

132. Jacomy M, Venturini T, Heymann S, Bastian M. 2014. ForceAtlas2, a continuous graph layout algorithm for handy network visualization designed for the Gephi software. PLoS One 9:e98679.

## REFERENCES File S1

1. Aguirre M, Collins MD. 1992. Phylogenetic analysis of *Alloiococcus otitis* gen. nov., sp. nov., an organism from human middle ear fluid. Int J Syst Bacteriol 42:79–83.

2. Yarza P, Richter M, Peplies J, Euzeby J, Amann R, Schleifer KH, Ludwig W, Glockner FO, Rossello-Mora R. 2008. The All-Species Living Tree project: a 16S rRNA-based phylogenetic tree of all sequenced type strains. Syst Appl Microbiol 31:241–50.

3. Yilmaz P, Parfrey LW, Yarza P, Gerken J, Pruesse E, Quast C, Schweer T, Peplies J, Ludwig W, Glockner FO. 2014. The SILVA and “All-species Living Tree Project (LTP)” taxonomic frameworks. Nucleic Acids Res 42:D643–8.

4. Escapa IF, Chen T, Huang Y, Gajare P, Dewhirst FE, Lemon KP. 2018. New Insights into Human Nostril Microbiome from the Expanded Human Oral Microbiome Database (eHOMD): a Resource for the Microbiome of the Human Aerodigestive Tract. mSystems 3.

5. Brugger SD, Eslami SM, Pettigrew MM, Escapa IF, Henke MT, Kong Y, Lemon KP. 2020. *Dolosigranulum pigrum* Cooperation and Competition in Human Nasal Microbiota. mSphere 5:e00852–20.

## REFERENCES File S2

1. Carattoli A, Zankari E, García-Fernández A, Voldby Larsen M, Lund O, Villa L, Møller Aarestrup F, Hasman H. 2014. In SilicoDetection and Typing of Plasmids using PlasmidFinder and Plasmid Multilocus Sequence Typing. Antimicrobial Agents and Chemotherapy 58:3895–3903.

2. Ma X, Li J, Xiong Y, Zhai Z, Ren F, Hao Y. 2016. Characterization of a Rolling-Circle Replication Plasmid pM411 from *Lactobacillus plantarum* 1–3. Current Microbiology 73:820–826.

3. Prechtl RM. 2019. Formation and structure of exopolysaccharides of meat starter cultures. Technical University of Munich.

4. McArthur AG, Waglechner N, Nizam F, Yan A, Azad MA, Baylay AJ, Bhullar K, Canova MJ, De Pascale G, Ejim L, Kalan L, King AM, Koteva K, Morar M, Mulvey MR, O’Brien JS, Pawlowski AC, Piddock LJV, Spanogiannopoulos P, Sutherland AD, Tang I, Taylor PL, Thaker M, Wang W, Yan M, Yu T, Wright GD. 2013. The Comprehensive Antibiotic Resistance Database. Antimicrobial Agents and Chemotherapy 57:3348–3357.

5. Jia B, Raphenya AR, Alcock B, Waglechner N, Guo P, Tsang KK, Lago BA, Dave BM, Pereira S, Sharma AN, Doshi S, Courtot M, Lo R, Williams LE, Frye JG, Elsayegh T, Sardar D, Westman EL, Pawlowski AC, Johnson TA, Brinkman FSL, Wright GD, McArthur AG. 2017. CARD 2017: expansion and model-centric curation of the comprehensive antibiotic resistance database. Nucleic Acids Research 45:D566–D573.

6. Ikebe T, Wada A, Inagaki Y, Sugama K, Suzuki R, Tanaka D, Tamaru A, Fujinaga Y, Abe Y, Shimizu Y, Watanabe H, Working Group for Group ASiJ. 2002. Dissemination of the phage-associated novel superantigen gene speL in recent invasive and noninvasive *Streptococcus pyogenes* M3/T3 isolates in Japan. Infect Immun 70:3227–33.

7. Beres SB, Sylva GL, Barbian KD, Lei B, Hoff JS, Mammarella ND, Liu M-Y, Smoot JC, Porcella SF, Parkins LD, Campbell DS, Smith TM, McCormick JK, Leung DYM, Schlievert PM, Musser JM. 2002. Genome sequence of a serotype M3 strain of group A *Streptococcus*: Phage-encoded toxins, the high-virulence phenotype, and clone emergence. Proceedings of the National Academy of Sciences 99:10078.

8. Yoon BH, Chang H-I. 2015. Genomic annotation for the temperate phage EFC-1, isolated from Enterococcus faecalis KBL101. 160:601–604.

9. Stanley E, Fitzgerald GF, Marrec CL, Fayard B, van Sinderen D. 1997. Sequence analysis and characterization of phi O1205, a temperate bacteriophage infecting *Streptococcus thermophilus* CNRZ1205. Microbiology (Reading) 143 (Pt 11):3417–3429.

10. Ma YL, Lu CP. 2008. Isolation and identification of a bacteriophage capable of infecting *Streptococcus suis* type 2 strains. Vet Microbiol 132:340–7.

11. Liu YP, Tang Q, Zhang JZ, Tian LF, Gao P, Yan XX. 2017. Structural basis underlying complex assembly and conformational transition of the type I R-M system. Proc Natl Acad Sci U S A 114:11151–11156.

12. Roberts RJ, Vincze T, Posfai J, Macelis D. 2015. REBASE--a database for DNA restriction and modification: enzymes, genes and genomes. Nucleic Acids Res 43:D298–9.

13. Xu SY, Corvaglia AR, Chan SH, Zheng Y, Linder P. 2011. A type IV modification-dependent restriction enzyme SauUSI from *Staphylococcus aureus* subsp. *aureus* USA300. Nucleic Acids Res 39:5597–610.

14. Sullivan MB, Huang KH, Ignacio-Espinoza JC, Berlin AM, Kelly L, Weigele PR, Defrancesco AS, Kern SE, Thompson LR, Young S, Yandava C, Fu R, Krastins B, Chase M, Sarracino D, Osburne MS, Henn MR, Chisholm SW. 2010. Genomic analysis of oceanic cyanobacterial myoviruses compared with T4-like myoviruses from diverse hosts and environments. 12:3035–3056.

15. Lu Z, Breidt F, Jr., Fleming HP, Altermann E, Klaenhammer TR. 2003. Isolation and characterization of a *Lactobacillus plantarum* bacteriophage, phiJL-1, from a cucumber fermentation. Int J Food Microbiol 84:225–35.

16. Edwards GB, Luna AJ, Hernandez AC, Kuty Everett GF. 2015. Complete Genome Sequence of *Citrobacter freundii* Myophage Moon. Genome Announcements 3:e01427–14.

17. Kauffman KM, Brown JM, Sharma RS, Vaninsberghe D, Elsherbini J, Polz M, Kelly L. 2018. Viruses of the Nahant Collection, characterization of 251 marine Vibrionaceae viruses. Scientific Data 5.

18. Smith BA, Leligdon C, Baltrus DA. 2019. Just the Two of Us? A Family of *Pseudomonas* Megaplasmids Offers a Rare Glimpse Into the Evolution of Large Mobile Elements. Genome Biology and Evolution doi:10.1093/gbe/evz066.

19. Janssen PJ, Van Houdt R, Moors H, Monsieurs P, Morin N, Michaux A, Benotmane MA, Leys N, Vallaeys T, Lapidus A, Monchy S, Médigue C, Taghavi S, McCorkle S, Dunn J, Van Der Lelie D, Mergeay M. 2010. The Complete Genome Sequence of *Cupriavidus metallidurans* Strain CH34, a Master Survivalist in Harsh and Anthropogenic Environments. PLoS ONE 5:e10433.

20. Del Rio B, Sánchez-Llana E, Redruello B, Magadan AH, Fernández M, Martin MC, Ladero V, Alvarez MA. 2019. *Enterococcus faecalis* Bacteriophage 156 Is an Effective Biotechnological Tool for Reducing the Presence of Tyramine and Putrescine in an Experimental Cheese Model. Frontiers in Microbiology 10.

21. Buttimer C, Hendrix H, Oliveira H, Casey A, Neve H, McAuliffe O, Ross RP, Hill C, Noben J-P, O’Mahony J, Lavigne R, Coffey A. 2017. Things Are Getting Hairy: Enterobacteria Bacteriophage vB_PcaM_CBB. 8.

22. Lopez MS, Hodde MK, Chamakura KR, Kuty Everett GF. 2014. Complete Genome of *Bacillus megaterium* Podophage Page. Genome Announcements 2:e00332–14.

23. Monod C, Repoila F, Kutateladze M, Tetart F, Krisch HM. 1997. The genome of the pseudo T-even bacteriophages, a diverse group that resembles T4. J Mol Biol 267:237–49.

24. Mehdizadeh Gohari I, Kropinski AM, Weese SJ, Parreira VR, Whitehead AE, Boerlin P, Prescott JF. 2016. Plasmid Characterization and Chromosome Analysis of Two netF+ Clostridium perfringens Isolates Associated with Foal and Canine Necrotizing Enteritis. PLOS ONE 11:e0148344.

25. Joensen KG, Scheutz F, Lund O, Hasman H, Kaas RS, Nielsen EM, Aarestrup FM. 2014. Real-time whole-genome sequencing for routine typing, surveillance, and outbreak detection of verotoxigenic Escherichia coli. J Clin Microbiol 52:1501–10.

